# Phylogeny and Genetic Diversity of Philippine Native Pigs (*Sus scrofa*) as Revealed by Mitochondrial DNA Analysis

**DOI:** 10.1101/2022.10.16.510730

**Authors:** Joy B. Banayo, Kathlyn Louise V. Manese, Agapita J. Salces, Takahiro Yamagata

**Affiliations:** Animal Genetics and Breeding, Graduate School of Bioagricultural Sciences, Nagoya University, Furocho, Chikusa, Nagoya 464-8601, Japan; Animal Breeding Division, Institute of Animal Science, College of Agriculture and Food Science, University of the Philippines Los Baños, College, Laguna 4031, Philippines

**Keywords:** Mitochondrial DNA d-loop control region, Indigenous pig, Livestock, Biodiversity, Haplotype, Conservation

## Abstract

Philippine native pigs (PhNP) are small black pigs domesticated in rural communities in the Philippines. They are valued locally for their various sociocultural roles. Recently, considerable literature has accumulated in the field of native pig production and marketing. However, there is limited research on the genetic diversity of PhNP. No previous study has investigated the evolutionary relatedness among native pigs from various islands and provinces in Luzon and Visayas, Philippines. In addition, a much-debated question is whether the PhNP were interbreeding with or even domesticated from endemic wild pigs. This study aims to clarify some of the uncertainties surrounding the identity and classification of PhNP based on mitochondrial DNA (mtDNA) signatures. Native pig samples (n=157) were collected from 10 provinces in Luzon and the Visayas, Philippines. Approximately 650 base pairs of the mtDNA d-loop control region were sequenced and analyzed together with publicly available sequences. Pairwise-distance analysis showed genetic separation of North and South Luzon (SL) and the clustering of SL with Visayan pigs. Phylogenetic analysis showed that the PhNP clustered within 3 recognized Asian pig domestication centers: D2 (East Asia), D7 (Southeast Asia) and the Cordillera clade (sister to the Lanyu). We identified 19 haplotypes (1-38 samples each), forming 4 haplogroups i.e. North Luzon, South Luzon and Visayas, Asian mix and the Cordillera cluster. No endemic wild pig mtDNA was detected in the native pig population, but the use of wild pig for interspecific hybridization was observed. This study showed that the Philippine native pigs have ancestral origin from at least 3 *Sus scrofa* lineage and that they were not domesticated from the endemic wild pigs of the Philippines.

## INTRODUCTION

The Philippine native pig (PhNP) is a small black pig domesticated in rural communities in the Philippines. They were introduced around 4000 years ago (Amano et al. 2013; Piper et al. 2009b) and are currently valued for their heat tolerance, disease resistance and meat quality (Santiago et al. 2016). As such, the Congress of the Philippines has passed a local legislation (Senate Bill 821 Philippine Native Animal Development Act of 2019) that promotes their conservation and development. This bill defines the native animals as “refer[s] to breeds of chickens, pigs, cattle, goats, sheep, ducks and other domesticated farm animals that are more adapted to the environmental conditions of the Philippines, having emerged through a long process of natural selection”. Therefore, the word ‘native’ was used as early as 1980 (Eusebio 1980; Maddul 1991; Peñalba 1993; Bondoc 1998; Bondoc 2008; Basilio 2016; Dichoso et al., 2022) pertaining to the adaptability of the animals to local conditions and their important sociocultural role, rather than its evolutionary context. True native and endemic pigs in the Philippines are referred to as wild pigs i.e. *Sus philippensis, S. cebifrons, S. oliveri, and S. ahoenobarbus* (Heaney et al. 2020; Ingicco et al. 2017; Lucchini et al. 2005; Groves 1997).

The utility of the native pig varies depending on the geographic location within the country. There are at least 7,641 islands (Larena et al. 2021; NAMRIA 2017, 2018) in the Philippines and at least 100 ethnic groups (Reyes et al. 2016) and 179 dialects (PSA 2022). In the highland provinces of the Cordillera region, native pigs are considered an important sociocultural component and are used in rituals and festivities (Lapeña and Acabado 2017). Various ethnic meat products, such as *etag* and *kiniing*, have been developed from this animal. Recently, there has been renewed interest in the native pig for the *lechon* (whole-roasted pig) delicacy in the mainstream market, for which native pigs are preferred over transboundary breeds due to the superior quality of their meat. Taken together, these developments suggest that the native pig can play a crucial role in contributing to food security and livelihood in rural communities.

Previous genetic studies have shown that native pigs from Batanes (Northern Philippines) (Li et al. 2017), South Luzon (Dichoso et al. 2022) and Visayas (Layos et al. 2022a, b) possess ancient mtDNA signatures initially found in the miniature Lanyu breed of Taiwan (Wu et al. 2007a). Layos et al. (2022b) further identified an A143T mutation that distinguishes the Philippine native from the Lanyu. Microsatellite analysis showed that they are distinct from Duroc and Yorkshire but maybe related to the Berkshire (Oh et al. 2014). Furthermore, farm productivity of several native pig populations showed subtle differences (DOST 2017). The current study will compare the genetic relatedness among pigs in Luzon, Visayas, and global breeds to support local efforts on native pig conservation (Philippine Native Animal Development Act of 2019; Santiago et al. 2016; Monleon 2005). In addition, we intend to clarify the much debated question on whether native pigs were interbreeding with or even domesticated from the endemic wild pigs in the Philippines (Eusebio 1980; Bondoc 1998; Bondoc 2008).

This study aims to clarify some of the uncertainties surrounding the ancestry and relatedness of native pigs in the Philippines. We will examine native pigs from 10 provinces using mtDNA signatures.

## MATERIALS AND METHODS

### Samples and Sampling Site

PhNP samples (ear notch tissue or hair follicle) were collected from 10 provinces in the Philippines (Fig. 1) following FAO (2011) guidelines on the use of unrelated individuals and unbiased sampling. Ear tissue was clipped from the ear and stored in ethanol at −20 °C until needed. This study included a total of 200 pig samples (n=157 native, n=43 transboundary breeds) collected from *(i)* smallhold farms in Benguet (n=15), Mt. Province (n=7), Kalinga, (n=32), Isabela (n=22), Nueva Viscaya (n=17), Quirino (n=3), Quezon (n=21), Marinduque (n=20), combined Eastern, Northern and Western Samar (n=18), Leyte (n=2), *(ii)* the National Swine and Poultry Research Development Center, Bureau of Animal Industry (BAI), Tiaong Stock Farm, Tiaong, Quezon for Berkshire (BAI BS; n=10), *(iii)* BAI-accredited swine breeding farms in the Philippines for purebred BS (n=3), Landrace (LR; n=10), Large White (LW; n=10) and Duroc (DC; n=10).

**Fig. 1.**
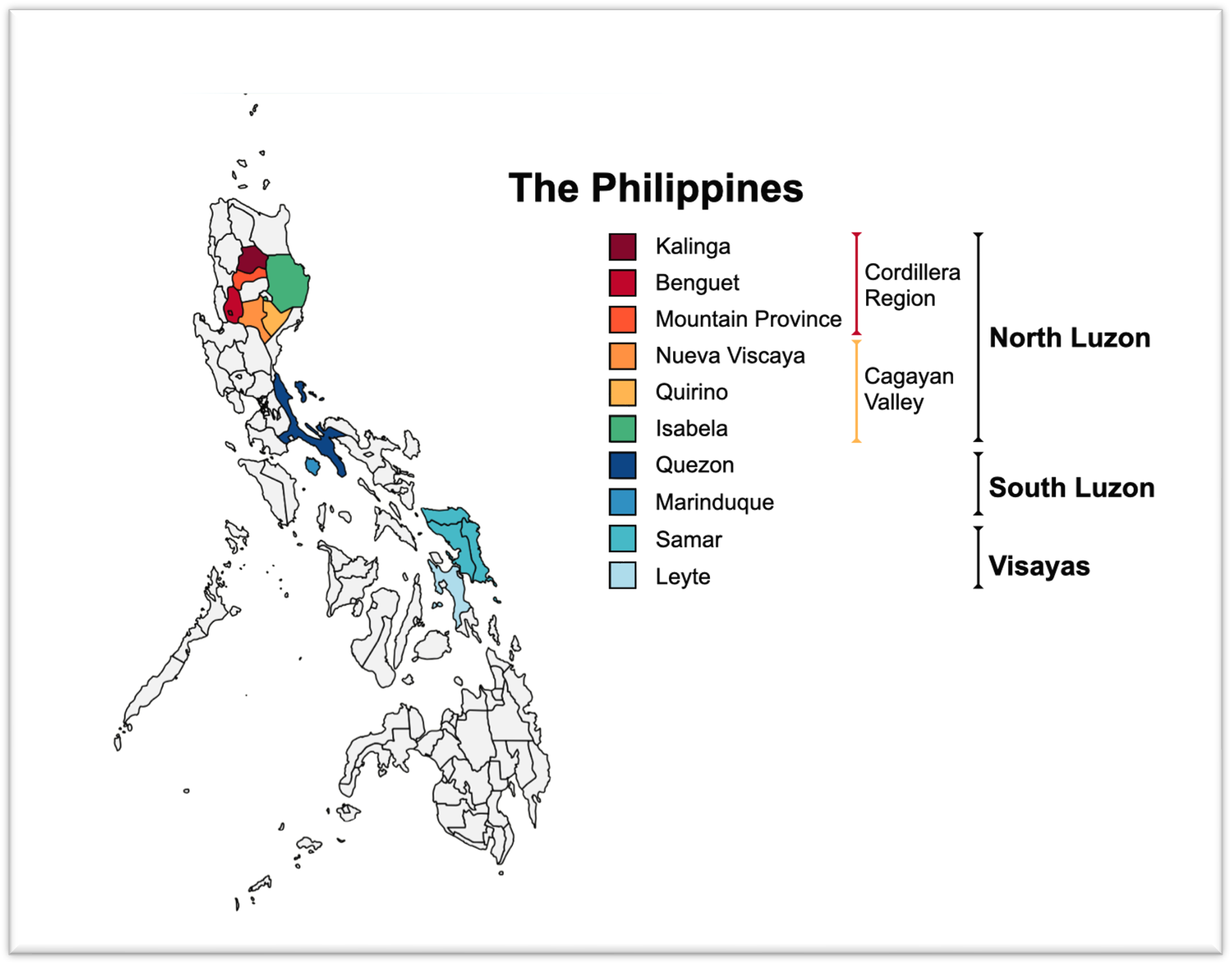
Sampling sites of the native pigs in the Philippines

### DNA Extraction and PCR Amplification

Ear notch samples (20 mg) or hair follicles (5 pieces) were rehydrated in phosphate-buffered saline and extracted using a GF-1 Tissue DNA Extraction Kit (Vivantis Technologies, Malaysia) following the manufacturer’s protocol. The partial mtDNA d-loop region (about 650 base pairs (bp)) was amplified using a single pair of primers: forward 5’-CAACCAAAACAAGCATTCCATTC-3’, reverse 5’-ATGAGTTCCATGAAGTCCAGC-3’. PCR was prepared in 25 μL reactions using 2x *Taq* Mastermix (Vivantis Technologies, Malaysia) containing a final concentration of 1 U *Taq* DNA polymerase, 1.5 mM MgCl_2_, 0.2 mM dNTPs, and 0.2 μM each of the forward and reverse primers and 20 ng DNA. The PCR products were sequenced directly by double pass standard sequencing with purification by a third-party service provider (Macrogen Inc., Seoul, South Korea). Nucleotide sequence data reported are available in the DDBJ/EMBL/GenBank databases under the accession numbers OM363266 – OM363454.

### Additional Sequences

Additional sequences were downloaded from the NCBI Nucleotide and RefSeq database (NCBI Resource Coordinators, 2016), including the following: Panay (MN625805-28, n=23, Layos et al. 2022b), Batanes (KP987306-7, n=2, Li et al. 2017), *S. cebifrons* (NC_023541.1, Si unpublished), and African warthog *Phacochoerus africanus* (NC_008830.1, Wu et al. 2007b), and domestic and wild pigs from Europe and Asia (Online Resource 1).

### mtDNA Genetic Diversity (p-Distances)

Genetic divergence was computed within groups and between groups in MEGA X v10.2.4 using the Tamura 3-parameter model with 1000 bootstrap replicates (Kumar et al. 2018). Sequences were first grouped into 6 Philippine subpopulations (n=19 to 50) and 4 transboundary breeds (DC n=26, BS n=17, LR n=31 and LW n=47). Samples were pooled to create the subpopulations Cordillera (Benguet, Mt. Province, Kalinga), Cagayan Valley (Isabela, Nueva Viscaya and Quirino), and Samar (Eastern, Western, Northern Samar, Leyte). The rate of variation within each site was modeled with a gamma distribution (shape parameter 0.39). All ambiguous positions were removed from each sequence pair (pairwise deletion). A total of length of 577 bp was used in the final dataset.

### Phylogenetic Analysis

Sequences (n=785 taxa, of which n=191 are from this study after successful sequencing, Online Resource 1) were aligned by MUSCLE in MEGA X software v10.2.4. DNA sequences were trimmed to equal length, and gaps were deleted (398 and 538 positions tested). Phylogeny was constructed by various methods, such as neighbor-joining (NJ), maximum likelihood (ML) and minimum evolution (ME), with 1000 bootstrap replicates each. The DNA substitution model of Tamura and Nei 1993 (TN93) was identified as best for the data based on the lowest BIC value in the Find Best DNA Model option in MEGA X. The substitution matrix was estimated using TN93 with assumptions of 5 gamma categories and heterogeneity patterns, where gaps were treated as complete deletions. A subset of 367 taxa was used to reconstruct a more refined tree using the Tamura 1993 model of DNA substitution with all sites used. The phylogenetic tree was rooted by *P. africanus*.

### Haplotype Analysis

Two haplotype networks were created using 672 taxa at 577 bp length and using a subset of 290 taxa at 538 bp length. Haplotype networks were constructed by median-joining using Population Analysis with Reticulate Tress (PopART) (Leigh and Bryant 2015; Bandelt et al. 1999) software. The Philippine haplotype sequence, number of haplotypes (h), number of polymorphic sites (S), haplotype diversity (Hd), and Tajima’s *D* were generated in DNA Sequence Polymorphism (DnaSP) v6.12.03 (Rozas et al. 2017) software using a length of 538 bp of 175 PhNP sequences (from this study and Genbank).

## RESULTS

### Genetic Divergence

The genetic divergence among the pig breeds was analyzed by pairwise distances. The native pigs were found to be diverged from DC (0.0167) and LR (0.0161), but may not be diverged from LW (0.0019) and BS (0.0016) (Table 1). Among the native pigs, Quezon was most diverged from BS (0.0059) and LW (0.0060). Native pigs closest (below 0.0030) to the BS were Cordillera, Cagayan, and Panay, and those to LW were Marinduque and Panay. Pairwise distances among transboundary breeds were 0.0001-0.0119, and among native pigs were 0.0001-0.0079. Among the native pigs, genetic distances of intermediate values (0.0055-0.0079) separated the North Luzon and South Luzon – Visayas populations. Short distances (≤ 0.0005) were observed between Cordillera and Cagayan Valley in North Luzon and among the islands of Marinduque, Samar and Panay of South Luzon – Visayas. Quezon pigs were relatively closer to Samar and Marinduque (0.0008-0.0012) and most distant from Panay (0.0016). Within-group distances ranged from 0.0058 (Marinduque) to 0.0140 (Cordillera) in natives and 0.0080 (DC) to 0.0120 (BS) in transboundary breeds.

**Table 1.** Estimates of genetic divergence by p-distances within and between populations of Philippine native and transboundary pigs

| Populations | Sub-Populations | Within Group (d) | Net Between Groups (d) |  |  |  |  |  |  |  |  |  |  |
| --- | --- | --- | --- | --- | --- | --- | --- | --- | --- | --- | --- | --- | --- |
|  | Philippine native (n=177) | 0.0130 ± 0.0029 | Ph native | North Luzon |  | South Luzon |  | Visayas |  | Transboundary |  |  |  |
|  |  |  |  | Cordillera | Cagayan Valley | Quezon | Marinduque | Samar | Panay | Duroc | Berkshire | Landrace | Large White |
| North Luzon | Cordillera (n=52) | 0.0140 ± 0.0037 | - | - |  |  |  |  |  |  |  |  |  |
|  | Cagayan Valley (n=43) | 0.0139 ± 0.0035 | - | 0.0003 | - |  |  |  |  |  |  |  |  |
| South Luzon | Quezon (n=19) | 0.0088 ± 0.0019 | - | 0.0070 | 0.0079 | - |  |  |  |  |  |  |  |
|  | Marinduque (n=19) | 0.0058 ± 0.0023 | - | 0.0055 | 0.0065 | 0.0012 | - |  |  |  |  |  |  |
| Visayas | Samar (n=20) | 0.0068 ± 0.0024 | - | 0.0058 | 0.0063 | 0.0008 | 0.0005 | - |  |  |  |  |  |
|  | Panay (n=24) | 0.0068 ± 0.0023 | - | 0.0052 | 0.0058 | 0.0016 | 0.0001 | 0.0002 | - |  |  |  |  |
| Transboundary | Duroc (n=36) | 0.0080 ± 0.0021 | 0.0167 ± 0.0045 | 0.0159 | 0.0153 | 0.0236 | 0.0210 | 0.0211 | 0.0203 | - |  |  |  |
|  | Berkshire (n=17) | 0.0120 ± 0.0029 | 0.0016 ± 0.0005 | 0.0029 | 0.0023 | 0.0059 | 0.0035 | 0.0034 | 0.0027 | 0.0110 | - |  |  |
|  | Landrace (n=31) | 0.0087 ± 0.0023 | 0.0161 ± 0.0046 | 0.0158 | 0.0152 | 0.0221 | 0.0200 | 0.0199 | 0.0194 | 0.0001 | 0.0102 | - |  |
|  | Large White (n=47) | 0.0111 ± 0.0029 | 0.0019 ± 0.0005 | 0.0037 | 0.0038 | 0.0060 | 0.0024 | 0.0033 | 0.0020 | 0.0119 | 0.0007 | 0.0117 | - |
| Overall mean diversity |  |  | 0.0163 ± 0.0034 |  |  |  |  |  |  |  |  |  |  |

### Phylogenetic Analysis

Phylogenetic analysis confirmed that the PhNP belong to the *S. scrofa* Asian clade (Fig. 2). They can be further divided into 6 subclades: Clade 1-5 and Cordillera clade and with a bootstrap support of 15, 10, 45, 55, 12 and 98%, respectively. Clade 1 included pigs from Quezon, Marinduque, Samar and Nicobari Island. Clade 1 also contained PhNP of D7 haplotype previously identified by Layos et al. (2022a, b). Clade 2 included 18 pigs from various provinces in North Luzon and mixed with Asian (Formosa wild boar and native pigs from China, Vietnam, Sri Lanka and India), BS and Yorkshire (YS) pigs. Clade 3-5 was composed of pigs from various provinces and mixed with LW/YS and BS, among others. Only the Cordillera clade has a distinct node (98% bootstrap) composed of 38 pigs from the Cordillera region (Kalinga, Benguet, Mt. Province), Nueva Viscaya, Isabela and Batanes Island. We reconstructed the phylogenetic tree using a shorter sequence (398 positions) and more taxa with the same result (data not shown).

**Fig. 2.**
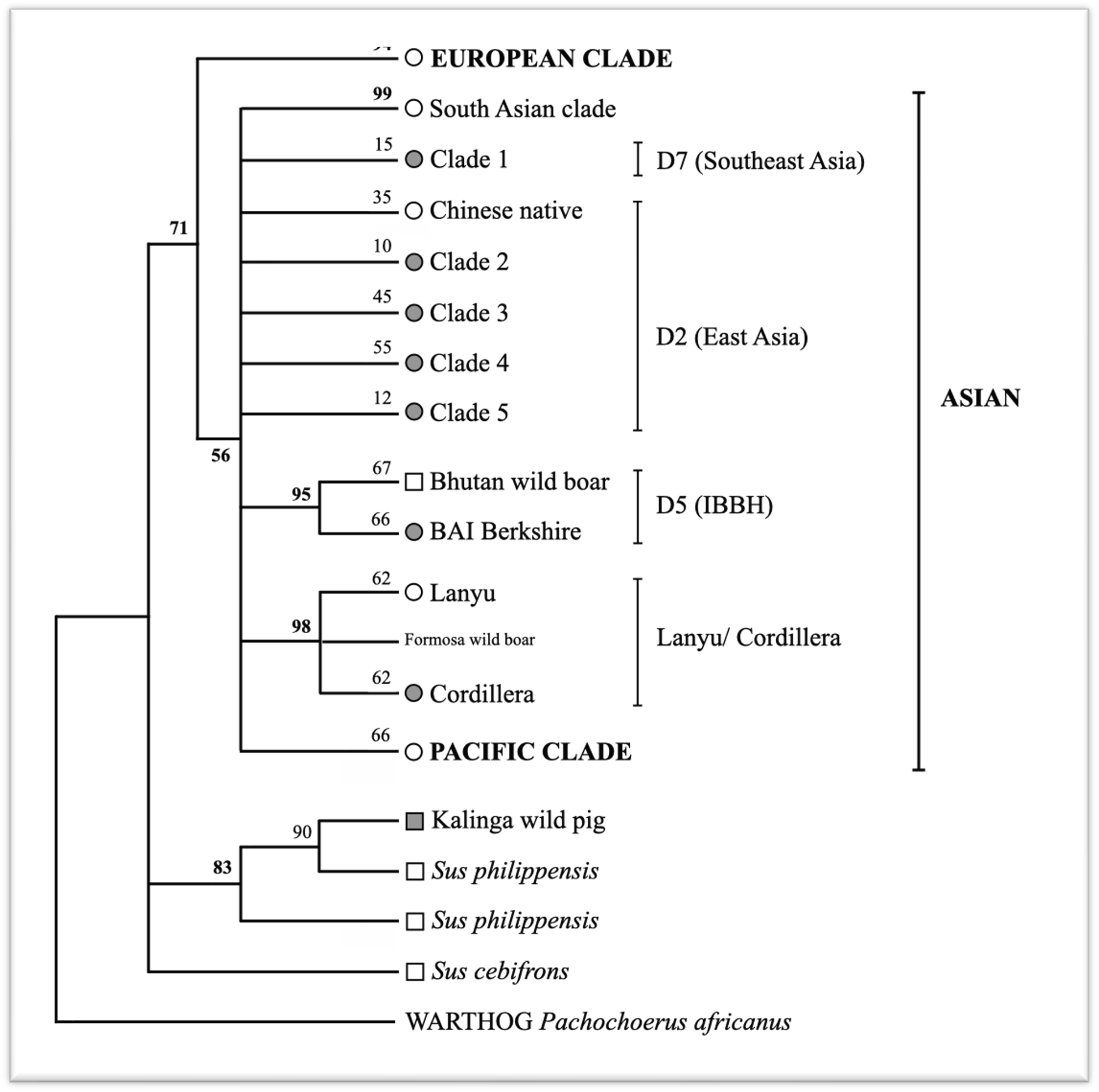
Phylogenetic tree of mtDNA signatures from Philippine native, Asian, European and wild pigs. Percentage bootstrap support is shown next to the branches. D2, D5, and D7 are indicated according to Larson et al. (2005) and Layos et al. (2022a). Node marker: circles-domestic pigs, squares-wild pig or wild boar, gray fill-Philippine samples collected for this study

On the other hand, our DC and LR samples clustered within the European clade (D1), while the LW and BS clustered within the general Asian clade (D2). However, BAI BS clustered distinctly with Bhutan wild boar such as HQ318431 belonging to Larson et al. D5 (2005) (Nidup unpublished) and is therefore not used further in the analysis. Furthermore, a single sample from Kalinga (OM363266) clustered with the *S. philippensis* clade with 90% bootstrap support (Fig. 2) and is therefore excluded from the Kalinga native pig population.

### Haplotype Analysis

Haplotype analysis of the PhNP revealed 19 haplotypes, Ph_1 to Ph_19 (Fig. 3 and 4). Ten haplotypes were unique to the Philippines and nine were shared with native and wild boars from Asia (Chinese, Jeju, Lanyu, Okinawa, Nicobari, Vietnamese, Ryukyu, Celebes wild pig) and Europe (Tamsworth, French wild boar, prehistoric Austrian) and with transboundary breeds (LW, DC, BS). The haplotype network revealed at least 4 haplogroups, hereby referred to as the Cordillera Cluster (CC), North Luzon Cluster (NLC), South Luzon-Visayas Cluster (SLVC) and Asian Mix Cluster (AMC) (Fig. 4). The number of haplotypes per clade was 2, 2, 8, and 7 for the CC (Ph_1-2), NLC (Ph_10-11 and 2 median vectors), SLVC (Ph_12-19 and 2 median vectors) and AMC (Ph_3-9), respectively. Each haplotype was represented by 1 (Ph_14) to 38 (Ph_1) samples. Ph_9 is shared between a large proportion of LW/YS and native pigs from various provinces. The DC, LR and LW samples in this study showed European haplotypes, while the BS showed an Asian haplotype (not shown). The haplotype distribution per region was 13, 9, and 11 in North Luzon, South Luzon and the Visayas, respectively (Fig. 5A). Each province had 1 (Leyte) to 8 (Samar) haplotypes (Fig. 5B). The details of the haplotype are presented in Online Resources 2 to 4.

**Fig. 3.**
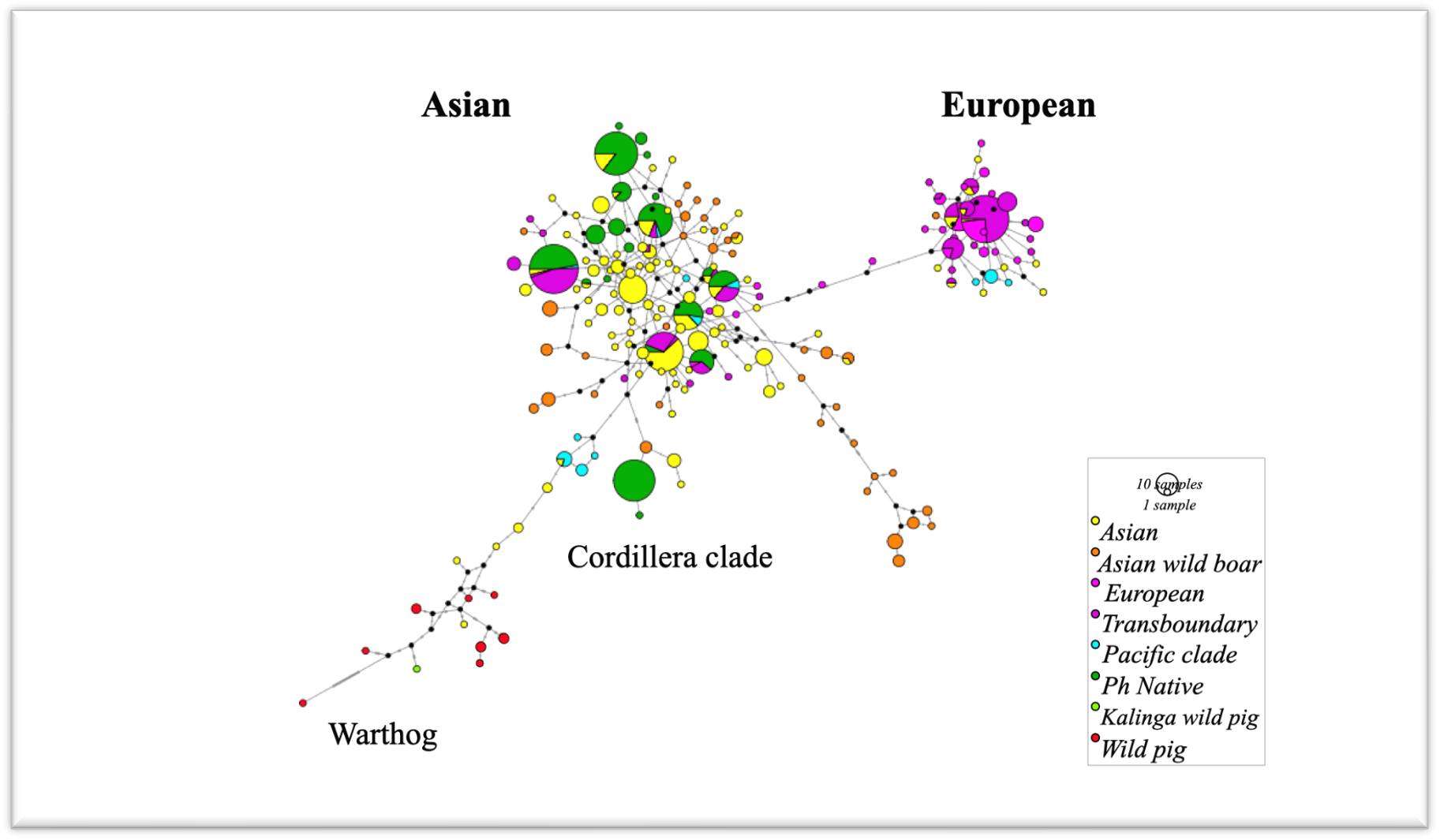
Haplotype network of mtDNA control region depicting the relationship of Philippine pigs to other breeds. The Philippine (Ph) native pigs have 19 haplotypes and belong to the general Asian clade. Number of sequences used: Asian n=208, Asian wild boar n=69, European n=54, Transboundary n=134, Pacific clade n=22, Ph Native n=178, Kalinga wild pig n=1, Wild pig n=6, total n=672

**Fig. 4.**
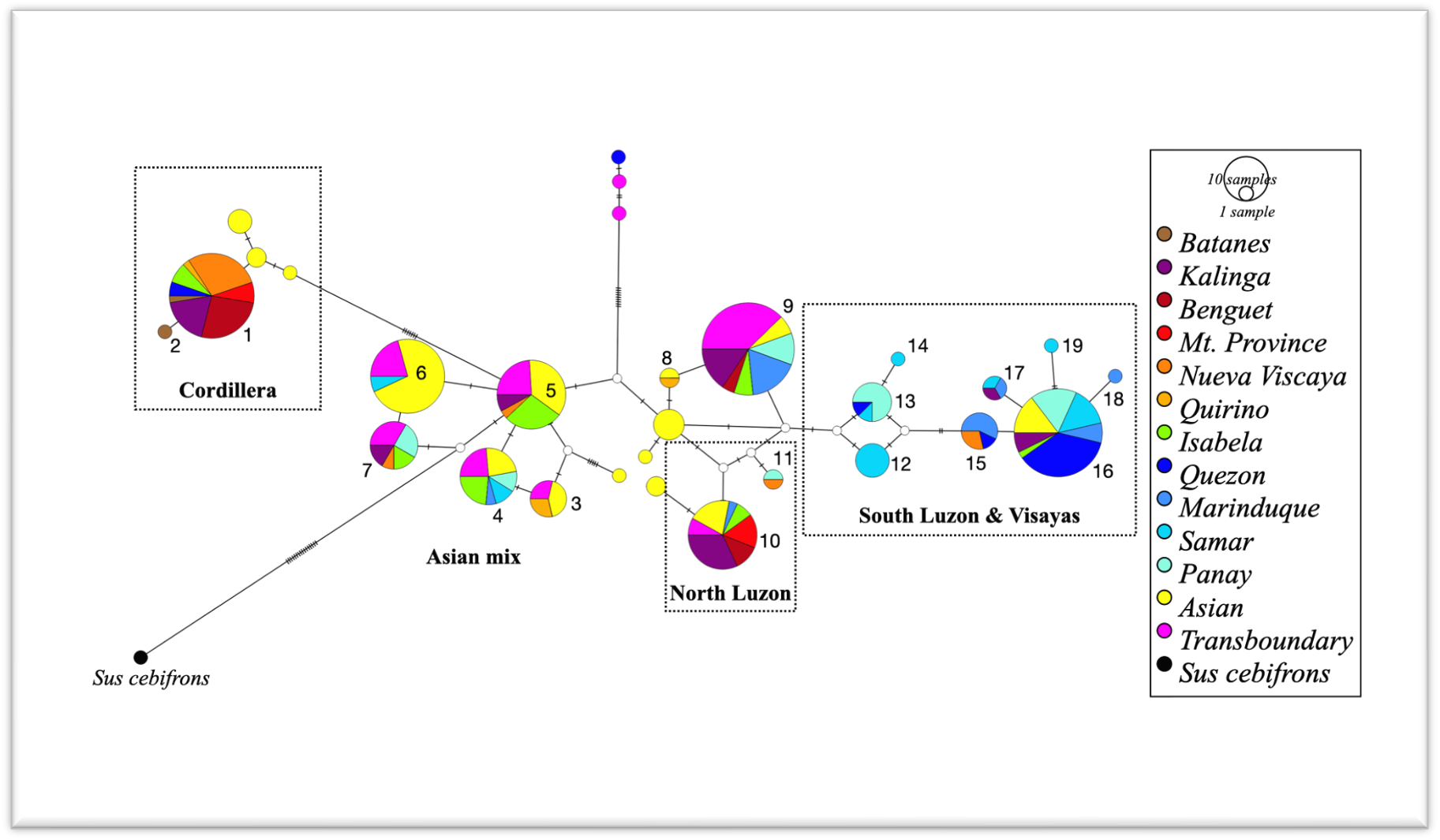
Haplotype network of mtDNA control region that is either unique or shared among Philippine native, Asian and transboundary pigs. Numbers indicate Philippine (Ph) haplotypes, which can be assigned to 4 haplogroups: (1) Cordillera, (2) North Luzon, (3) South Luzon & Visayas, and (4) all remaining, to the general Asian mix clade. Samples used: Ph n=180, Asian n=67, transboundary breeds n=43. Hatch marks denote single mutations. White circles are median vectors showing consensus sequences between haplotypes representing potentially unsampled or extinct relatives

**Fig. 5.**
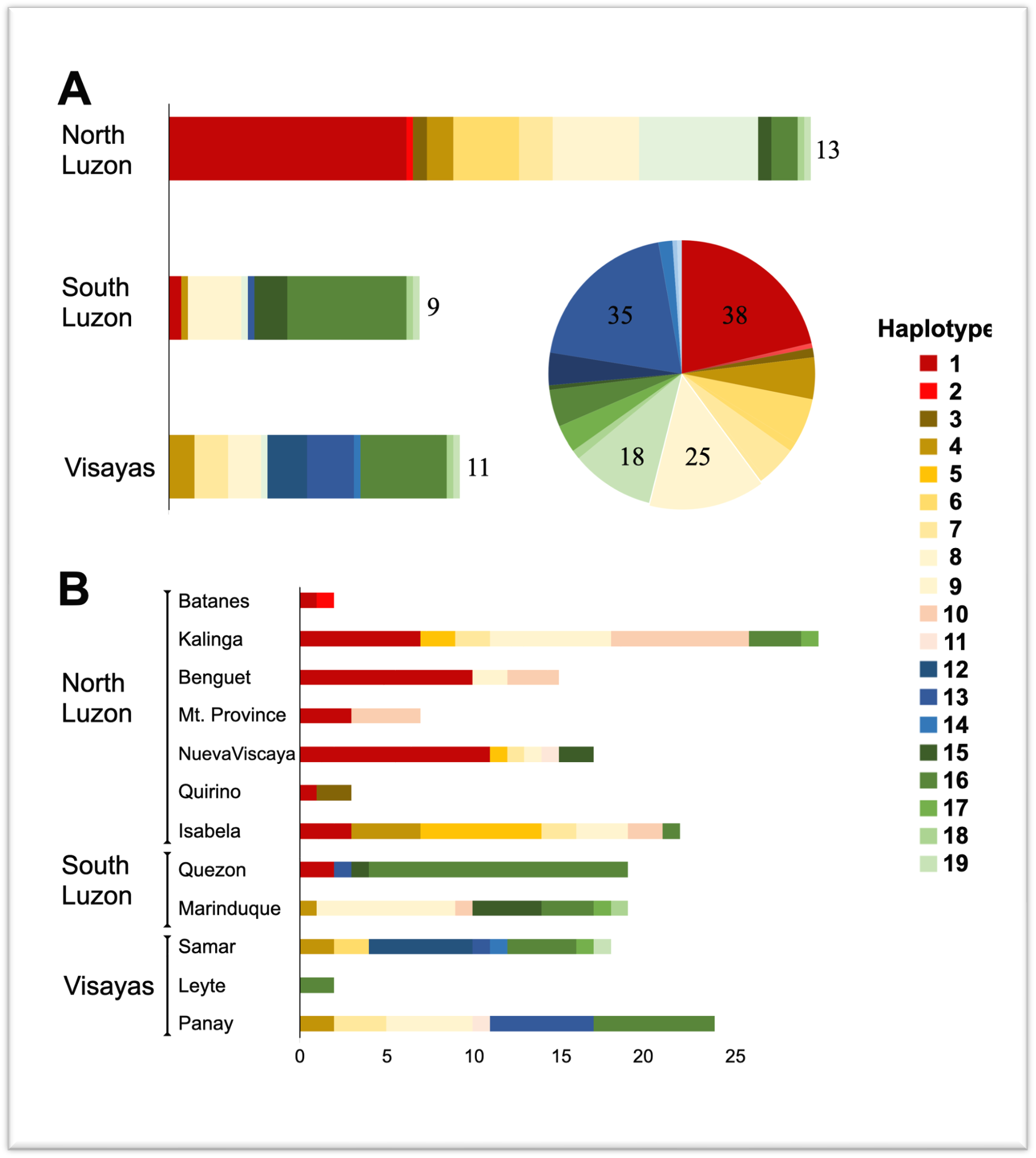
Distribution of the 19 haplotypes per Philippine region (A) and province (B). The size of the bar and pie graph is proportional to the sampling size, the number of haplotypes for each region is shown on the side of the bar graph and the number of sample size of major haplotypes is shown in the pie chart (A).

## DISCUSSION

The PhNP showed a considerable genetic distance from DC and LR but were closer to LW and BS (Table 1). Microsatellite analysis by Oh et al. (2014) showed similar results. It must be noted that LW and BS are known to have Asian mtDNA due to their intensive upgrading with Asian pigs (Giuffra et al. 2000). The genetic distances between native pig populations correlated with their geographic distance, suggesting some isolation between pigs from North Luzon and those in South Luzon and the Visayas (Fig. 1). The same trend of NL vs. SL separation was also observed among Philippine native cattle using BoLA gene markers (Takeshima et al. 2014). The higher mtDNA diversity observed in the Cordillera and Cagayan Valley (encompassing 4 Provinces in North Luzon) was due to the presence of multiple haplotypes (n=13) from the 2 varying lineages (Asian and Cordillera clade) (Fig. 2).

There are at least 7 pig domestication centers recognized and denoted as D1-D7 as well as other cryptic domestication sites (Larson et al. 2005; Tanaka et al. 2008; Layos et al. 2022a). Phylogenetic analysis showed that PhNP clustered to various lineages such as the ancient Cordillera/Lanyu clade and the general Asian Clade (D2 and D7) (Fig. 2). Pigs from NL clustered in the Cordillera clade or in the D2 (East Asian) clade, while pigs from SLV clustered within the D7 (Southeast Asian). This study provides additional evidence on the multiple ancestry of the PhNP as observed by previous studies (Layos et al., 2022a,b; Dichoso et al. 2022).

Of the 19 haplotypes, the Ph_1 haplotype was the major haplotype in this study (Fig. 4). The median-joining network showed that the Ph_1 haplotype was 10 mutations and 2 median vectors away from the nearest Ph haplotype (Ph_5). It was, however, only one mutation away from the Formosa wild boar with-Lanyu-sign-lineage (KP9897300). Similarly, the type I Lanyu (EF375877.3) is also characterized by a single mutation from the Formosa wild boar but at another position (Fig. 3 and 4). Layos identified a single mutation (A143T transversion) that distinguishes the PhNP from the Lanyu (Layos et al 2022a). Our data suggest that the Cordillera clade is sister to the Lanyu and hint at the potential domestication of a common ancestral wild boar in the Philippines, whether it is the Formosa or another yet undiscovered wild boar. There were speculations of an ancient land connection between [Mountain Province] Cordillera and Formosa during early Tertiary which facilitated the dispersal of several Himalayan plants into both areas at the same time and through the same channels (Merrill 1926). Central Cordillera region is believed to have originated as a series of small islands about 30 million years ago (MYA) that formed what is now the Central Cordillera (Heaney 2016). Therefore, the geological history of the Cordillera region is too old to discount the natural dispersal of the pig belonging to the Cordillera and the Lanyu clade. Currently, the Cordillera is one of the most diverse and important biotas in the Philippines with many endemic genera (Heaney 2016; Heaney et al. 2005). The Lanyu was estimated to have diverged at 0.6 MYA earlier than other East Asian wild boars (Li et al. 2017). The CC haplotype was the major mtDNA signature of PhNP in Benguet, Mt. Province, Kalinga and Nueva Viscaya. These provinces are connected by the Cordillera and Caraballo Mountain Range, where indigenous communities prefer this type of pig. The CC haplotype continues into Isabela, the Palawan region and Bohol (Layos et al. 2022b), possibly by human-mediated translocation. Ancient mtDNA haplotypes were also found in Philippine native chickens (Thomson et al. 2014). We show long-term genetic continuity between early and modern domestic pigs as was observed in China (Larson et al. 2010).

In the Philippines, crosses of LW, LR and DC were common in smallhold production (Peñalba 2005). These are still the main breeds used locally (DOST 2018). The shared haplotype (Ph_9) of PhNP and LW/YS may indicate localized crossbreeding for the production of *lechon* pig. Crossbreeding with transboundary breeds have replaced the mtDNA signatures of several Asian native pigs (Charoensook et al. 2019; Zhang et al. 2018; Zhang et al. 2016). On the other hand, the shared haplotype with BS could be due to shared mtDNA ancestry, since the BS possess Asian type mtDNA (Giuffra et al. 2000). It is interesting that the BS is most distant from Quezon native indicating that the *Berkjala* crossbreed was unable to contribute BS mtDNA in the modern population of South Luzon. The *Bekjala* was developed in the early 1900s using BS and Laguna native pig (Matias 1995; Eusebio 1980) but is presumably extinct (Arboleda et al. 2003).

Being traditionally free-ranged, the PhNP were hypothesized to be interbreeding with or even domesticated from Philippine endemic wild pigs (Eusebio 1890; Bondoc 1998; Bondoc 2008). We confirm only one pig, described as wild by the owner, to be an *S. philippensis* type (Fig. 2). No wild pig mtDNA was detected in pigs described as native, corroborating the *S. scrofa* ancestry of the PhNP. Since mtDNA is only maternally inherited, the depth of hybridizations and genetic introgressions between native and endemic wild cannot be ascertained in this study. Even the wild boar mtDNA were often not detectable in modern domestic pigs (Tanaka et al. 2008; Larson et al. 2005; Cho et al. 2009; Watanobe et al. 1999). These cases prove the limitation of the mtDNA in detecting hybridizations and paternal lineage (Sasaki and Sato 2021; Moore 1995). Wild boar Y-chromosome analysis did not show clear correlation with mtDNA clades (Choi et al. 2020). Furthermore, Frantz et al. (2013) showed that gene flow was higher between domestic *S. scrofa* breeds than between domestic and wild relatives. In the Philippines, however, interspecific hybridization is a major threat to the genetic integrity of the wild pig populations (Heaney et al. 2020; Meijard and Melleti 2018; Melleti et al. 2018; Tabaranza et al. 2018).

We provide evidence on the local preference for interspecific hybridizations in Kalinga, Philippines. The close cultural relationship of the Filipino people with both wild and native pigs alike have occurred since ancient times (Piper et al. 2009a). The Philippine native cattle have similar interspecific origins (Aquino et al. 2006). Interspecific and intergeneric admixtures (observed between pygmy hog and the wild boar) have been proven to be important biological driving forces for adaptation (Liu et al. 2019; Ai et al. 2015). The adaptability of the PhNP in low input systems could be influenced by interspecific hybridization with endemic wild pigs. Furthermore, we observed that farms intentionally keep warty pigs to mate with PhNP suggesting that interspecific hybrids are advantaged in low input farming. This poses a challenge on the co-management of native and wild pigs to address both the need for adaptability in the former and for genetic integrity of the latter. Our data provide additional evidence on the expanse and hybridization of *S. philippensis* in Kalinga province (Meijard and Melleti 2018; Heaney et al 2005).

## CONCLUSION

This study set out to examine the genetic relatedness of the PhNP to clarify their identity and classification. The mtDNA of PhNP did not show close affinity to that of the endemic wild pigs in the Philippines, indicating that the PhNP had other genetic origins. Further analysis, revealed multiple ancestral origins from East and Southeast Asia. We observed local preference for interspecific hybridization potentially to improve the adaptability of PhNP. This study contributes to our understanding of Asian domestic pig ancestry, hybridization and dispersal.

## Supporting information

Online Resource 1-5

## ACKNOWLEDGMENTS

We acknowledge the support of the Department of Science and Technology - Philippine Council for Agriculture, Aquatic and Natural Resources Research and Development (DOST-PCAARRD), Philippines, for sample collection through the native pig program. We further acknowledge local collaborators at Kalinga State University, Benguet State University, Isabela State University, Nueva Viscaya State University, National Swine and Poultry Research and Development Center, Marinduque State College and Eastern Samar State University (Online Resource 5) for their assistance with the sampling. Last, we thank Dr. Elpidio M. Agbisit, Jr. for providing the purebred swine DNA.

## STATEMENTS AND DECLARATIONS

### Author Contributions

All authors contributed to the study conception and design. Material preparation and data collection were performed by Joy B. Banayo and Kathlyn Louise V. Manese. Data analysis and drafting and revising of the manuscript were performed by Joy B. Banayo and Takahiro Yamagata. All authors commented on previous versions of the manuscript. All authors read and approved the final manuscript.

### Data Availability

Nucleotide sequence data reported are available in the DDBJ/EMBL/GenBank databases under the accession numbers OM363266 - OM363454. All data will be available on reasonable request.

### Ethics approval

The sampling protocol described in this study was reviewed and approved by the University of the Philippines Los Baños Institutional Animal Care and Use Committee (24 June 2021/UPLB-2021-025).

## REFERENCES

Ai H, Fang X, Yang B, Huang Z, Chen H, Mao L, Zhang F, Zhang L, Cui L, He W, Yang J, Yao X, Zhou L, Han L, Li J, Sun S, Xie X, Lai B, Su Y, Lu Y, Yang H, Huang T, Deng W, Nielsen R, Ren J, Huang L (2015) Adaptation and possible ancient interspecies introgression in pigs identified by whole-genome sequencing. Nat Genet 47(3):217–225. https://doi.org/10.1038/ng.3199

Arboleda CR, Ranola RF, Bondoc OL, Lambio AL, Peñalba FF, Castro NL, Lanuza MB, Flores AB (2003) Policy studies on animal genetic improvement and production in the Philippines: I. Review of Significant Legislative Actions Since 1900. Philippine Journal of Veterinary and Animal Sciences 29(1):1–15. Retrieved from http://www.ejournals.ph/form/cite.php?id=8684

Aquino GMB, Laude RP, Jianlin H, Sevilla CS, Hanotte O (2006) Genetic structure of selected cattle populations in the Philippines using microsatellites. Philippine J Vet Anim Sci 32(1):1–9

Bandelt HJ, Forster P, Röhl A (1999) Median-joining networks for inferring intraspecific phylogenies. Mol Biol Evol 16(1):37–48. https://doi.org/10.1093/oxfordjournals.molbev.a026036

Basilio EB Jr. (2016) Genetic diversity of Ifugao and Kalinga native pigs (Sus scrofa L.) using morphometrics and molecular markers. [Unpublished doctoral thesis]. University of the Philippines Los Baños. 94 pp

Bondoc OL (1998) Biodiversity of livestock and poultry genetic resources in the Philippines. University of the Philippines Los Baños and the Department of Science and Technology, Philippines

Bondoc OL (2008) Animal breeding: Principles and practice in the Philippine context. The University of the Philippines Press, Quezon City, Philippines

Charoensook R, Gatphayak K, Brenig B, Knorr C (2019) Genetic diversity analysis of Thai indigenous pig population using microsatellite markers. Asian-Australas J Anim Sci 32(10):1491–1500. https://doi.org/10.5713/ajas.18.0832

Cho IC, Han SH, Fang M, Lee SS, Ko MS, Lee H, Lim HT, Yoo CK, Lee JH, Jeon JT (2009) The robust phylogeny of Korean wild boar (Sus scrofa coreanus) using partial D-loop sequence of mtDNA. Mol Cells 28(5):423–430. https://doi.org/10.1007/s10059-009-0139-3

Choi SK, Kim KS, Ranyuk M, Babaev E, Voloshina I, Bayarlkhagva D, Chong JR, Ishiguro N, Yu L, Min MS, Lee H, Markov N (2020) Asia-wide phylogeography of wild boar (Sus scrofa) based on mitochondrial DNA and Y-chromosome: Revising the migration routes of wild boar in Asia. PLoS ONE 15(8):e0238049. https://doi.org/10.1371/journal.pone.0238049

Dichoso MS, Cruz RJD, Vega RSA, Manuel MCC, Li KY, Ju YT, Laude RP (2022) Phylogenetic analysis of Philippine native pigs (Sus scrofa L.) from Batanes, Quezon, and Marinduque based on mitochondrial DNA d-loop markers. Philippine J Sci 151(3):1267–1276

DOST (2017) Philippine native pig breed information system. Philippine Council for Agriculture, Aquatic and Natural Resources Research and Development, Department of Science and Technology (DOST), Los Baños, Laguna, Philippines. https://pab-is.pcaarrd.dost.gov.ph/nativepigs. Accessed 15 July 2021

DOST (2018) SwineCart. Philippine Council for Agriculture, Aquatic and Natural Resources Research and Development, Department of Science and Technology (DOST), Los Baños, Laguna, Philippines. https://swinecart.work/customer/public-products. Accessed 110 January 2022

Eusebio JA (1980) Pig production in the tropics (Intermediate tropical agriculture series). Longman Group Ltd., Harlow, Essex, United Kingdom. 128 pp.

FAO (2011) Molecular genetic characterization of animal genetic resources. FAO Animal Production and Health Guidelines. No. 9. Rome.

Frantz LA, Schraiber JG, Madsen O, Megens HJ, Bosse M, Paudel Y, Semiadi G, Meijaard E, Li N, Crooijmans RP, Archibald AL, Slatkin M, Schook LB, Larson G, Groenen MA (2013) Genome sequencing reveals fine scale diversification and reticulation history during speciation in Sus. Genome Biol 14(9):R107. https://doi.org/10.1186/gb-2013-14-9-r107

Giuffra E, Kijas JMH, Amarger V, Carlborg O, Jeon JT, Andersson L (2000) The origin of the domestic pig: independent domestication and subsequent introgression. Genetics 154(4):1785–1791

Groves CP (1997) Taxonomy of the wild pigs (Sus) in the Philippines. Zoological J Linne Soc 120:163–191

Heaney LR, Duya MRM, Ong PS, Ingle ANR, Reginaldo AA, Yaptinchay AA, Mildenstein T, Balete D, Sedlock J, Carino AB, Alviola P (2020) Mammals. P.21-36. In Philippine red list of threatened wild fauna: Part 1-Vertebrates. (van de Ven, ed). Biodiversity Management Bureau, Department of Environment and Natural Resources (DENR), Quezon City, Philippines. 116 pp.

Heaney LR, Balete DS, Gee GV, Tabao MVL, Rickart EA, Tabaranza BRJ (2005) Preliminary report on the mammals of Balbasang, Kalinga Province, Luzon. Sylvatrop: The Philippine Forest Research Journal 13:51–62.

Ingicco T, Piper PJ, Amano N, Paz VJ, Pawlik AF (2017) Biometric differentiation of wild Philippine pigs from introduced Sus scrofa in modern and archaeological assemblages. Int J Osteoarchaeol 27(5):768–784. https://doi.org/10.1002/oa.2592

Kumar S, Stecher G, Li M, Knyaz C, Tamura K (2018) MEGA X: Molecular evolutionary genetics analysis across computing platforms. Mol Biol Evol 35(6):1547–1549. https://doi.org/10.1093/molbev/msy096

Lapeña QG, Acabado SG (2017) Resistance through rituals: The role of Philippine “native pig” (Sus scrofa) in Ifugao feasting and socio-political organization. J Archaeol Sci Rep 13:583–594. https://doi.org/10.1016/j.jasrep.2017.05.009

Larena M, Sanchez-Quinto F, Sjödin P et al. (2021) Multiple migrations to the Philippines during the last 50,000 years. Proc Natl Acad Sci USA 118(13):e2026132118. https://doi.org/10.1073/pnas.2026132118

Larson G, Dobney K, Albarella U, Fang M, Matisoo-Smith E, Robins J, Lowden S, Finlayson H, Brand T, Willerslev E, Rowley-Conwy P, Andersson L, Cooper A (2005) Worldwide phylogeography of wild boar reveals multiple centers of pig domestication. Science 307(5715):1618–1621. https://doi.org/10.1126/science.1106927

Larson G, Liu R, Zhao X, Yuan J, Fuller D, Barton L, Dobney K, Fan Q, Gu Z, Liu XH, Luo Y, Lv P, Andersson L, Li N (2010) Patterns of East Asian pig domestication, migration, and turnover revealed by modern and ancient DNA. Proc Natl Acad Sci USA 107(17):7686–7691. https://doi.org/10.1073/pnas.0912264107

Layos JKN, Geromo RB, Espina DM, Nishibori M (2022a) Insights on the historical biogeography of Philippine domestic pigs and its relationship with continental domestic pigs and wild boars. PloS ONE 17(3):e0254299. https://doi.org/10.1371/journal.pone.0254299

Layos JKN, Godinez CJP, Liao LM, Yamamoto Y, Masangkay JS, Mannen H, Nishibori M (2022b) Origin and demographic history of Philippine pigs inferred from mitochondrial DNA. Front Genet 12:8233641. https://doi.org/10.3389/fgene.2021.823364

Leigh JW, Bryant D (2015) POPART: Full-feature software for haplotype network construction. Methods Ecol Evol 6(9):1110–1116. https://doi.org/10.1111/2041-210X.12410

Li KY, Li KT, Yang CH, Hwang MH, Chang SW, Lin SM, Wu HJ, Basilio EB Jr, Vega RSA, Laude RP, Ju YT (2017) Insular East Asia pig dispersal and vicariance inferred from Asian wild boar genetic evidence. J Anim Sci 95(4):1451–1466. https://doi.org/10.2527/jas.2016.1117

Liu L, Bosse M, Megens HJ, Frantz LAF, Lee YL, Irving-Pease EK, Narayan G, Groenen MAM, Madsen O (2019) Genomic analysis on pygmy hog reveals extensive interbreeding during wild boar expansion. Nat Commun 10(1):1992. https://doi.org/10.1038/s41467-019-10017-2

Lucchini V, Meijaard E, Diong CH, Groves CP, Randi E (2005) New phylogenetic perspectives among species of South-east Asian wild pig (Sus sp.) based on mtDNA sequences and morphometric data. J Zoology 266(1):25–35. https://doi.org/10.1017/S0952836905006588

Maddul S (1991) Production, management and characteristics of the native pigs in the Cordillera. [Unpublished doctoral thesis]. University of the Philippines Los Baños. 120 pp.

Matias JM (1995) Conservation systems of animal genetic resources in the Philippines. In: The Proceedings of the 3^rd^ MAFF International Workshop on Genetic Resources: Efficient Conservation and Effective Use, 5-7 December, NIAR, Tsukuba, Japan

Meijaard E, Melletti M (2018) Philippine warty pig Sus philippensis (Nehring, 1886). pp. 157–161. in Ecology, Evolution and Management of Wild Pigs and Peccaries: Implications for Conservation. (M. Melletti and E. Meijaard, eds.). Cambridge University Press, United Kingdom. 480 pp.

Melletti M, Meijard E, Przybylska L (2018) Visayan warty pig Sus cebifrons (Heude, 1888). pp. 150–156. in Ecology, Evolution and Management of Wild Pigs and Peccaries: Implications for Conservation. (M. Melletti and E. Meijaard, eds.). Cambridge University Press, United Kingdom. 480 pp.

Monleon AM (2005) Local conservation efforts for the Philippine native pig (Sus domesticus) in Marinduque. Philippine J Vet and Anim Sci 32(1):79–86

Moore WS (1995) Inferring phylogenies from mtDNA variation: mitochondrial-gene trees versus nuclear gene trees. Evolution 49(4):718–726. https://doi.org/10.1111/j.1558-5646.1995.tb02308.x

NAMRIA, 2017. News and Events: Administrator Tiangco welcomes (2017) National Mapping and Resource Information Authority, Department of Environment and Natural Resources (NAMRIA-DENR), Philippines. Available online at http://www.namria.gov.ph

NAMRIA, 2018. Annual Report: Building a geospatially-empowered Philippines. National Mapping and Resource Information Authority, Department of Environment and Natural Resources (NAMRIA-DENR), Philippines. p.7. Available online at httP://www.namria.gov.ph

NCBI Resource Coordinators (2016) Database resources of the National Center for Biotechnology Information. Nucleic Acids Res 44(D1), D7–D19. https://doi.org/10.1093/nar/gkv1290

Oh JD, Cacho RCC, Choi JY, Seo JH, Song KD, Vega RSA, Santiago RC, Octura JER, Kim SW, Kim CW, Kim SH, Kong SH, Lee HK, Cho BW (2014) Genetic analysis of Philippine native pigs (Sus scrofa L.) using microsatellite loci. Philippine J Sci 143(1): 89–96

Peñalba FF (2005) Backcrossing of Landrace and Large White crossbred as an alternative system in the production of female breeding pigs for the smallholders. Agricultural Science and Technology Information (AGRIS) Database at http://www.agris.fao.org. Accessed 15 November 2021

Philippine Native Animal Development Act of 2019 (2019). 18^th^ Congress of the Republic of the Philippines. Retrieved from the Legislative Digital Resources at https://issuances-library.senate.gov.ph/bills/senate-bill-no-821-18th-congress-republic

Piper PJ, Campos FZ, Hung H (2009a) A study of the animal bone recovered from pits 9 and 10 at the site of Nagsabaran in Northern Luzon, Philippines. Hukay 14:47–90

Piper PJ, Hung HC, Campos FZ, Bellwood P, and Santiago R (2009b) A 4000-year-old introduction of domestic pigs into the Philippine Archipelago: implications for understanding routes of human migration through Island Southeast Asia and Wallacea. Antiquity 83, 687–695. https://doi.org/10.1017/S0003598X00098914

PSA (2022) Methodology of data collection, processing, dissemination and analysis: Ethnicity and Language/Dialect Generally Spoken at Home. 2010 Census of Population and Housing. Philippines Statistics Office (PSA). Available online at http://www.psa.gov.ph. Accessed 22 July 2022

Reyes CM, Mina CD, Asis RD (2016) Inequality patterns among ethnic groups in the Philippines. 13th National Convention on Statistics, 3-4 October. Available online at http://www.psa.gov.ph. Accessed 22 July 2022

Rozas J, Ferrer-Mata A, Sánchez-DelBarrio JC, Guirao-Rico S, Librado P, Ramos-Onsins SE, Sánchez-Gracia A (2017) DnaSP 6: DNA sequence polymorphism analysis of large datasets. Mol Biol Evol 34: 3299–3302. https://doi.org/10.1093/molbev/msx248

Santiago RC, Lambio AL, Dimaranan KC (2016) Philippine native animals, source of pride and wealth, worth conserving and utilizing. National Swine and Poultry Research and Development Center (NSPRDC), Bureau of Animal Industry, Department of Agriculture, Quezon City. 200pp.

Sasaki T, Sato M (2021) Degradation of paternal mitochondria via mitophagy. Biochim Biophys Acta. 1865(6):129886. https://doi.org/10.1016/j.bbagen.2021.129886

Tabaranza DGE, Schütz E, Gonzalez JCT, Afuang LTM (2018) Mindoro warty pig Sus oliveri (Groves, 1997). pp. 162–169. in Ecology, Evolution and Management of Wild Pigs and Peccaries: Implications for Conservation. (M. Melletti and E. Meijaard, eds.). Cambridge University Press, United Kingdom. 480 pp.

Takeshima SN, Miyasaka T, Polat M, Kikuya M, Matsumoto Y, Mingala CN, Villanueva MA, Salces AJ, Onuma M, Aida Y (2014) The great diversity of major histocompatibility complex class II genes in Philippine native cattle. Meta Gene 2:176–190. https://www.doi.org/10.1016/j.mgene.2013.12.005

Tanaka K, Iwaki Y, Takizawa T, Dorji T, Tshering G, Kurosawa Y, Maeda Y, Mannen H, Nomura K, Dang VB, Chhum-phith L, Bouahom B, Yamamoto Y, Daing T, Namikawa T (2008) Mitochondrial diversity of native pigs in the mainland South and South-east Asian countries and its relationships between local wild boars. Anim Sci J 79(4):417–434. https://www.doi.org/10.1111/j.1740-0929.2008.00546.x

Thomson VA, Lebrasseur O, Austin JJ, Hunt TL, Burney DA, Denham T, Rawlence NJ, Wood JR, Gongora J, Girdland Flink L, Linderholm A, Dobney K, Larson G, Cooper A (2014) Using ancient DNA to study the origins and dispersal of ancestral Polynesian chickens across the Pacific. Proc Natl Acad Sci USA 111(13):4826–4831. https://doi.org/10.1073/pnas.1320412111

Zhang J, Jiao T, Zhao S (2016) Genetic diversity in the mitochondrial DNA D-loop region of global swine (Sus scrofa) populations. Biochem Biophys Res Commun. 473(4):814–820. https://doi.org/10.1016/j.bbrc.2016.03.125

Zhang J, Yang B, Wen X, Sun G (2018) Genetic variation and relationships in the mitochondrial DNA D-loop region of Qinghai indigenous and commercial pig breeds. Cell Mol Biol Lett 23:31. https://doi.org/10.1186/s11658-018-0097-x

Watanobe T, Okumura N, Ishiguro N, Nakano M, Matsui A, Sahara M, Komatsu M (1999) Genetic relationship and distribution of the Japanese wild boar (Sus scrofa leucomystax) and Ryukyu wild boar (Sus scrofa riukiuanus) analysed by mitochondrial DNA. Mol Ecol 8(9):1509–1512. https://doi.org/10.1046/j.1365-294x.1999.00729.x

Wu CY, Jiang YN, Chu HP, Li SH, Wang Y, Li YH, Chang Y, Ju YT (2007a) The type I Lanyu pig has a maternal genetic lineage distinct from Asian and European pigs. Anim Genet 38(5):499–505. https://doi.org/10.1111/j.1365-2052.2007.01646.x. PMID: 17894564

Wu GS, Yao YG, Qu KX, Ding ZL, Li H, Palanichamy MG, Duan ZY, Li N, Chen YS, and Zhang YP (2007b) Population phylogenomic analysis of mitochondrial DNA in wild boars and domestic pigs revealed multiple domestication events in East Asia. Genome Biol 8(11):R245. https://doi.org/10.1186/gb-2007-8-11-r245

