## Supplementary material for "Phylogeny and Genetic Diversity of Philippine Native Pigs (*Sus scrofa*) as Revealed by Mitochondrial DNA Analysis": Online Resource 1-5

**Online Resource 1** Pig samples used in the construction of the haplotype network

| **GENBANK ID** | **TRAIT** | **BREED** |
| --- | --- | --- |
| HQ318295.1 | Asian | Bhutan village pig |
| HQ318301.1 | Asian | Bhutan village pig |
| HQ318303.1 | Asian | Bhutan village pig |
| HQ318309.1 | Asian | Bhutan village pig |
| HQ318310.1 | Asian | Bhutan village pig |
| HQ318319.1 | Asian | Bhutan village pig |
| DQ779287.1 | Asian | Chinese domestic |
| DQ779353.1 | Asian | Chinese domestic |
| DQ779420.1 | Asian | Chinese domestic |
| AY230818.1 | Asian | Erhualian |
| AY230824.1 | Asian | Erhualian |
| AY230825.1 | Asian | Erhualian |
| AY230826.1 | Asian | Erhualian |
| KY964861.1 | Asian | Harbin White |
| KY964862.1 | Asian | Harbin White |
| KY964863.1 | Asian | Harbin White |
| KY964864.1 | Asian | Harbin White |
| KY964865.1 | Asian | Harbin White |
| KY964866.1 | Asian | Harbin White |
| KY964867.1 | Asian | Harbin White |
| KY964860.1 | Asian | Harbin White |
| KM016444.1 | Asian | Indian domestic |
| KM016445.1 | Asian | Indian domestic |
| KM016446.1 | Asian | Indian domestic |
| KM016447.1 | Asian | Indian domestic |
| KM016448.1 | Asian | Indian domestic |
| DQ779336.1 | Asian | Indonesian domestic |
| DQ779338.1 | Asian | Indonesian domestic |
| DQ841948.1 | Asian | Indonesian domestic |
| DQ841949.1 | Asian | Indonesian domestic |
| AY879785.1 | Asian | Jeju Native |
| AY879786.1 | Asian | Jeju Native |
| AY879787.1 | Asian | Jeju Native |
| AY879788.1 | Asian | Jeju Native |
| AY879789.1 | Asian | Jeju Native |
| AY879790.1 | Asian | Jeju Native |
| AY879791.1 | Asian | Jeju Native |
| AY879792.1 | Asian | Jeju Native |
| AY879793.1 | Asian | Jeju Native |
| DQ191229.1 | Asian | Jeju Native |
| DQ191230.1 | Asian | Jeju Native |
| DQ377716.1 | Asian | Jeju Native |
| DQ377717.1 | Asian | Jeju Native |
| DQ377718.1 | Asian | Jeju Native |
| AB041476.1 | Asian | Jinhua |
| JX068065.1 | Asian | Jinhua |
| JX068066.1 | Asian | Jinhua |
| JX068067.1 | Asian | Jinhua |
| KC469586.1 | Asian | Jinhua |
| GQ141898.1 | Asian | Jinyang White |
| GQ141899.1 | Asian | Jinyang White |
| DQ779421.1 | Asian | Korean Domestic |
| DQ779422.1 | Asian | Korean Domestic |
| DQ779524.1 | Asian | Korean Domestic |
| DQ779525.1 | Asian | Korean Domestic |
| DQ779526.1 | Asian | Korean Domestic |
| DQ779527.1 | Asian | Korean Domestic |
| DQ379166.2 | Asian | Laiwu |
| DQ379167.2 | Asian | Laiwu |
| DQ379168.2 | Asian | Laiwu |
| KR049158.1 | Asian | Laiwu |
| KR049159.1 | Asian | Laiwu |
| DQ518915.2 | Asian | Lanyu |
| DQ972936.3 | Asian | Lanyu |
| EF375877.3 | Asian | Lanyu |
| EU008084.1 | Asian | Lanyu |
| EU008085.1 | Asian | Lanyu |
| EU008086.1 | Asian | Lanyu |
| EU008087.1 | Asian | Lanyu |
| KP987303.1 | Asian | Lanyu |
| KP987304.1 | Asian | Lanyu |
| KP987305.1 | Asian | Lanyu |
| KY964873.1 | Asian | Liangshan |
| KY964874.1 | Asian | Liangshan |
| KY964875.1 | Asian | Liangshan |
| KY964876.1 | Asian | Liangshan |
| KY964877.1 | Asian | Liangshan |
| DQ779337.1 | Asian | Malaysian domestic |
| GQ141892.1 | Asian | Mashen |
| AB041474.1 | Asian | Meishan |
| AM040648.1 | Asian | Meishan |
| AM040649.1 | Asian | Meishan |
| AM040650.1 | Asian | Meishan |
| AM040651.1 | Asian | Meishan |
| AM040652.1 | Asian | Meishan |
| AY230821.1 | Asian | Meishan |
| AY230827.1 | Asian | Meishan |
| DQ379160.2 | Asian | Meishan |
| DQ379161.2 | Asian | Meishan |
| DQ379162.2 | Asian | Meishan |
| GQ169776.1 | Asian | Meishan |
| DQ379171.2 | Asian | Min |
| DQ379172.2 | Asian | Min |
| DQ379173.2 | Asian | Min |
| AB041469.1 | Asian | Miyazaki |
| DQ779411.1 | Asian | Myanmar domestic |
| DQ379132.2 | Asian | Neijiang |
| DQ379142.2 | Asian | Neijiang |
| DQ379143.2 | Asian | Neijiang |
| JX068068.1 | Asian | Neijiang |
| JX068069.1 | Asian | Neijiang |
| JX068070.1 | Asian | Neijiang |
| HQ318440.1 | Asian | Nepal village pig |
| HQ318441.1 | Asian | Nepal village pig |
| HQ318442.1 | Asian | Nepal village pig |
| HQ318443.1 | Asian | Nepal village pig |
| HQ318444.1 | Asian | Nepal village pig |
| HQ318445.1 | Asian | Nepal village pig |
| HQ318446.1 | Asian | Nepal village pig |
| HQ318447.1 | Asian | Nepal village pig |
| HQ318448.1 | Asian | Nepal village pig |
| HQ318449.1 | Asian | Nepal village pig |
| MT024618.1 | Asian | Nicobari |
| MT024619.1 | Asian | Nicobari |
| MT024620.1 | Asian | Nicobari |
| MT024621.1 | Asian | Nicobari |
| MT024622.1 | Asian | Nicobari |
| MT024623.1 | Asian | Nicobari |
| MT024624.1 | Asian | Nicobari |
| MT024625.1 | Asian | Nicobari |
| MT024626.1 | Asian | Nicobari |
| MT024627.1 | Asian | Nicobari |
| AB015092.1 | Asian | Okinawa Native |
| JX068437.1 | Asian | Pengzhou |
| JX068438.1 | Asian | Pengzhou |
| JX068439.1 | Asian | Pengzhou |
| KJ746664.1 | Asian | Penzhou |
| KY964883.1 | Asian | Qingyu |
| KY964884.1 | Asian | Qingyu |
| KY964885.1 | Asian | Qingyu |
| KY964886.1 | Asian | Qingyu |
| KY964887.1 | Asian | Qingyu |
| DQ379112.2 | Asian | Quanbei |
| DQ379135.2 | Asian | Quanbei |
| DQ379174.2 | Asian | Quanbei |
| KY964892.1 | Asian | Rongchang |
| KY964893.1 | Asian | Rongchang |
| KY964894.1 | Asian | Rongchang |
| KY964895.1 | Asian | Rongchang |
| KY964896.1 | Asian | Rongchang |
| KY964897.1 | Asian | Rongchang |
| DQ152884.2 | Asian | Shanggao |
| DQ152885.2 | Asian | Shanggao |
| DQ379186.2 | Asian | Shanggao |
| DQ152877.2 | Asian | Shengxian spotted |
| DQ379152.2 | Asian | Shengxian spotted |
| MF038765.1 | Asian | Shenxian |
| MF038766.1 | Asian | Shenxian |
| MF038767.1 | Asian | Shenxian |
| DQ779414.1 | Asian | Sri Lanka domestic |
| HQ318480.1 | Asian | Sri Lanka village pig |
| HQ318481.1 | Asian | Sri Lanka village pig |
| HQ318482.1 | Asian | Sri Lanka village pig |
| HQ318483.1 | Asian | Sri Lanka village pig |
| HQ318484.1 | Asian | Sri Lanka village pig |
| HQ318485.1 | Asian | Sri Lanka village pig |
| HQ318486.1 | Asian | Sri Lanka village pig |
| HQ318487.1 | Asian | Sri Lanka village pig |
| HQ318488.1 | Asian | Sri Lanka village pig |
| HQ318489.1 | Asian | Sri Lanka village pig |
| NC_014692 | Asian | *Sus scrofa taiwanensis* |
| DQ779417.1 | Asian | Taiwanese domestic |
| DQ779418.1 | Asian | Taiwanese domestic |
| GQ169775.1 | Asian | Taoyuan |
| AM778827.1 | Asian | Thailand native |
| AM779905.1 | Asian | Thailand native |
| JX068235.1 | Asian | Tibetan |
| JX068236.1 | Asian | Tibetan |
| KY964913.1 | Asian | Tibetan |
| KY964914.1 | Asian | Tibetan |
| DQ152875.2 | Asian | Tongcheng |
| DQ379138.2 | Asian | Tongcheng |
| DQ379147.2 | Asian | Tongcheng |
| DQ779432.1 | Asian | Vietnamese domestic |
| DQ779433.1 | Asian | Vietnamese domestic |
| DQ779434.1 | Asian | Vietnamese domestic |
| DQ779435.1 | Asian | Vietnamese domestic |
| DQ779436.1 | Asian | Vietnamese domestic |
| DQ779437.1 | Asian | Vietnamese domestic |
| KX982638.1 | Asian | Vietnamese domestic |
| KX982639.1 | Asian | Vietnamese domestic |
| KX982645.1 | Asian | Vietnamese domestic |
| KX982649.1 | Asian | Vietnamese domestic |
| KX982650.1 | Asian | Vietnamese domestic |
| JX068483.1 | Asian | Wujin |
| JX068484.1 | Asian | Wujin |
| JX068485.1 | Asian | Wujin |
| DQ379116.2 | Asian | Xiang |
| DQ379139.2 | Asian | Xiang |
| DQ379201.2 | Asian | Xiang |
| DQ152876.2 | Asian | Xiangxi |
| DQ152894.2 | Asian | Xiangxi |
| DQ152895.2 | Asian | Xiangxi |
| JX068486.1 | Asian | Yanan |
| JX068487.1 | Asian | Yanan |
| JX068488.1 | Asian | Yanan |
| DQ152883.2 | Asian | Yimeng |
| DQ379154.2 | Asian | Yimeng |
| DQ379175.2 | Asian | Yimeng |
| KR049166.1 | Asian | Yimeng |
| KR049167.1 | Asian | Yimeng |
| AB015093.1 | Asian | Yucatan Mini |
| DQ152871.2 | Asian | Yushan |
| DQ152878.2 | Asian | Yushan |
| DQ379141.2 | Asian | Yushan |
| DQ152874.2 | Asian | Zang |
| DQ152892.2 | Asian | Zang |
| DQ379102.2 | Asian | Zang |
| HQ318428.1 | Asian_wild_boar | Bhutan wild boar |
| HQ318429.1 | Asian_wild_boar | Bhutan wild boar |
| HQ318430.1 | Asian_wild_boar | Bhutan wild boar |
| HQ318431.1 | Asian_wild_boar | Bhutan wild boar |
| HQ318432.1 | Asian_wild_boar | Bhutan wild boar |
| HQ318433.1 | Asian_wild_boar | Bhutan wild boar |
| HQ318434.1 | Asian_wild_boar | Bhutan wild boar |
| HQ318435.1 | Asian_wild_boar | Bhutan wild boar |
| HQ318436.1 | Asian_wild_boar | Bhutan wild boar |
| HQ318437.1 | Asian_wild_boar | Bhutan wild boar |
| HQ318438.1 | Asian_wild_boar | Bhutan wild boar |
| HQ318439.1 | Asian_wild_boar | Bhutan wild boar |
| DQ779397.1 | Asian_wild_boar | Celebes Wild, *Sus celebensis* |
| DQ779398.1 | Asian_wild_boar | Celebes Wild, *Sus celebensis* |
| HM026640.1 | Asian_wild_boar | Chinese wild boar |
| JX068440.1 | Asian_wild_boar | Chinese wild boar |
| JX068441.1 | Asian_wild_boar | Chinese wild boar |
| JX068442.1 | Asian_wild_boar | Chinese wild boar |
| JX068443.1 | Asian_wild_boar | Chinese wild boar |
| JX068444.1 | Asian_wild_boar | Chinese wild boar |
| KP987287.1 | Asian_wild_boar | Formosan wild boar |
| KP987288.1 | Asian_wild_boar | Formosan wild boar |
| KP987289.1 | Asian_wild_boar | Formosan wild boar |
| KP987290.1 | Asian_wild_boar | Formosan wild boar |
| KP987291.1 | Asian_wild_boar | Formosan wild boar |
| KP987292.1 | Asian_wild_boar | Formosan wild boar |
| KP987293.1 | Asian_wild_boar | Formosan wild boar |
| KP987294.1 | Asian_wild_boar | Formosan wild boar |
| KP987295.1 | Asian_wild_boar | Formosan wild boar |
| KP987296.1 | Asian_wild_boar | Formosan wild boar |
| KP987297.1 | Asian_wild_boar | Formosan wild boar |
| KP987298.1 | Asian_wild_boar | Formosan wild boar |
| KP987299.1 | Asian_wild_boar | Formosan wild boar |
| KP987300.1 | Asian_wild_boar | Formosan wild boar |
| KP987301.1 | Asian_wild_boar | Formosan wild boar |
| KP987302.1 | Asian_wild_boar | Formosan wild boar |
| GQ141901.1 | Asian_wild_boar | Hybrid Liaoning wild boar |
| GQ141902.1 | Asian_wild_boar | Hybrid Liaoning wild boar |
| KM016430.1 | Asian_wild_boar | Indian wild boar |
| KM016431.1 | Asian_wild_boar | Indian wild boar |
| KM016432.1 | Asian_wild_boar | Indian wild boar |
| KM016433.1 | Asian_wild_boar | Indian wild boar |
| KM016434.1 | Asian_wild_boar | Indian wild boar |
| AY879777.1 | Asian_wild_boar | Korean wild boar |
| AY879778.1 | Asian_wild_boar | Korean wild boar |
| AY879779.1 | Asian_wild_boar | Korean wild boar |
| EF533692.1 | Asian_wild_boar | Korean wild boar |
| EF533693.1 | Asian_wild_boar | Korean wild boar |
| KY911743.1 | Asian_wild_boar | Korean wild boar |
| KY911744.1 | Asian_wild_boar | Korean wild boar |
| KY911745.1 | Asian_wild_boar | Korean wild boar |
| GQ141900.1 | Asian_wild_boar | Liaoning Wild |
| AY884620.1 | Asian_wild_boar | Nepal village pig |
| AB015088.1 | Asian_wild_boar | Ryukyu Wild |
| HQ318503.1 | Asian_wild_boar | Sri Lanka wild boar |
| HQ318504.1 | Asian_wild_boar | Sri Lanka wild boar |
| HQ318505.1 | Asian_wild_boar | Sri Lanka wild boar |
| HQ318506.1 | Asian_wild_boar | Sri Lanka wild boar |
| HQ318507.1 | Asian_wild_boar | Sri Lanka wild boar |
| HQ318508.1 | Asian_wild_boar | Sri Lanka wild boar |
| HQ318509.1 | Asian_wild_boar | Sri Lanka wild boar |
| HQ318510.1 | Asian_wild_boar | Sri Lanka wild boar |
| HQ318511.1 | Asian_wild_boar | Sri Lanka wild boar |
| HQ318512.1 | Asian_wild_boar | Sri Lanka wild boar |
| HQ318513.1 | Asian_wild_boar | Sri Lanka wild boar |
| HQ318514.1 | Asian_wild_boar | Sri Lanka wild boar |
| HQ318515.1 | Asian_wild_boar | Sri Lanka wild boar |
| HQ318516.1 | Asian_wild_boar | Sri Lanka wild boar |
| HQ318517.1 | Asian_wild_boar | Sri Lanka wild boar |
| HM026608.1 | European | Czechian wild boar |
| AB059651.1 | European | European wild boar |
| DQ379250.2 | European | French wild boar |
| HM026611.1 | European | French wild boar |
| AY230823.1 | European | German Landrace |
| AB041488.1 | European | Hampshire |
| DQ379016.2 | European | Hampshire |
| DQ379018.2 | European | Hampshire |
| DQ379037.2 | European | Hampshire |
| DQ379038.2 | European | Hampshire |
| HM026610.1 | European | Hungarian Wild |
| AY232862.1 | European | Iberian |
| AY232863.1 | European | Iberian |
| AY232864.1 | European | Iberian |
| CP071572.1 | European | Italian NS |
| HM026613.1 | European | Italian wild boar |
| DQ379021.2 | European | Large Black |
| DQ379022.2 | European | Large Black |
| DQ379118.2 | European | Large Black |
| DQ152859.2 | European | Manchado de Jabugo |
| DQ379063.2 | European | Mangalica |
| JX310707.1 | European | Mangalitsa |
| ZDQ379019.2 | European | Middle White |
| DQ152850.2 | European | Negro Canario |
| DQ379079.2 | European | Negro Iberico |
| AB041489.1 | European | Pietrain |
| AY230820.1 | European | Pietrain |
| AY232886.1 | European | Pietrain |
| AY232887.1 | European | Pietrain |
| DQ379099.2 | European | Pietrain |
| DQ379100.2 | European | Pietrain |
| DQ379229.2 | European | Pietrain |
| DQ379230.2 | European | Pietrain |
| DQ379231.2 | European | Pietrain |
| GQ141904.1 | European | Pietrain |
| GQ141905.1 | European | Pietrain |
| JX546549.1 | European | Pietrain |
| JX546550.1 | European | Pietrain |
| JX546551.1 | European | Pietrain |
| JX546552.1 | European | Pietrain |
| JX546553.1 | European | Pietrain |
| HM026612.1 | European | Polish Wild |
| MG92639 | European | prehistoric Austrian domestic |
| DQ152844.2 | European | Retinto |
| DQ152856.2 | European | Retinto |
| DQ152867.2 | European | Retinto |
| HM026614.1 | European | Romanian Wild |
| AY230819.1 | European | Schwabisch-Hallisches |
| HM026618.1 | European | Slovenia Wild |
| AY232868.1 | European | Spanish Wild |
| HM026607.1 | European | Spanish Wild |
| AY232890.1 | European | Spotted Black Jabugo |
| HM026615.1 | European | Swedish wild boar |
| DQ152896.2 | European | Tamworth |
| OM363266 | Kalinga wild pig | Philippine warty pig |
| DQ779528.1 | Pacific_Clade | Cook Islands pig |
| DQ994625.1 | Pacific_Clade | Cook Islands pig |
| DQ779429.1 | Pacific_Clade | French Polynesia pig |
| DQ779430.1 | Pacific_Clade | French Polynesia pig |
| DQ994637.1 | Pacific_Clade | Kapia |
| DQ994638.1 | Pacific_Clade | Kapia |
| DQ994618.1 | Pacific_Clade | Narave |
| DQ994632.1 | Pacific_Clade | Narave |
| HQ318519.1 | Pacific_Clade | Papua New Guinea pig |
| HQ318520.1 | Pacific_Clade | Papua New Guinea pig |
| HQ318521.1 | Pacific_Clade | Papua New Guinea pig |
| HQ318522.1 | Pacific_Clade | Papua New Guinea pig |
| HQ318523.1 | Pacific_Clade | Papua New Guinea pig |
| HQ318524.1 | Pacific_Clade | Papua New Guinea pig |
| HQ318581.1 | Pacific_Clade | Papua New Guinea pig |
| HQ318582.1 | Pacific_Clade | Papua New Guinea pig |
| HQ318583.1 | Pacific_Clade | Papua New Guinea pig |
| HQ318584.1 | Pacific_Clade | Papua New Guinea pig |
| DQ779530.1 | Pacific_Clade | Solomon Islands pig |
| AY884673.1 | Pacific_Clade | *Sus scrofa papuensis* |
| DQ994635.1 | Pacific_Clade | Vanuatu domestic |
| DQ994636.1 | Pacific_Clade | Vanuatu domestic |
| KP987306.1 | Ph_Native | Philippine native/ Batanes |
| KP987307.1 | Ph_Native | Philippine native/ Batanes |
| KP987308.1 | Ph_Native | Philippine native/ Batanes |
| OM363303 | Ph_Native | Philippine native/ Benguet |
| OM363304 | Ph_Native | Philippine native/ Benguet |
| OM363305 | Ph_Native | Philippine native/ Benguet |
| OM363306 | Ph_Native | Philippine native/ Benguet |
| OM363307 | Ph_Native | Philippine native/ Benguet |
| OM363308 | Ph_Native | Philippine native/ Benguet |
| OM363309 | Ph_Native | Philippine native/ Benguet |
| OM363310 | Ph_Native | Philippine native/ Benguet |
| OM363311 | Ph_Native | Philippine native/ Benguet |
| OM363312 | Ph_Native | Philippine native/ Benguet |
| OM363313 | Ph_Native | Philippine native/ Benguet |
| OM363314 | Ph_Native | Philippine native/ Benguet |
| OM363315 | Ph_Native | Philippine native/ Benguet |
| OM363316 | Ph_Native | Philippine native/ Benguet |
| OM363317 | Ph_Native | Philippine native/ Benguet |
| OM363318 | Ph_Native | Philippine native/ Benguet |
| OM363319 | Ph_Native | Philippine native/ Benguet |
| OM363320 | Ph_Native | Philippine native/ Benguet |
| OM363321 | Ph_Native | Philippine native/ Benguet |
| OM363322 | Ph_Native | Philippine native/ Benguet |
| OM363323 | Ph_Native | Philippine native/ Benguet |
| OM363324 | Ph_Native | Philippine native/ Benguet |
| OM363345 | Ph_Native | Philippine native/ Isabela |
| OM363346 | Ph_Native | Philippine native/ Isabela |
| OM363347 | Ph_Native | Philippine native/ Isabela |
| OM363348 | Ph_Native | Philippine native/ Isabela |
| OM363349 | Ph_Native | Philippine native/ Isabela |
| OM363350 | Ph_Native | Philippine native/ Isabela |
| OM363351 | Ph_Native | Philippine native/ Isabela |
| OM363352 | Ph_Native | Philippine native/ Isabela |
| OM363353 | Ph_Native | Philippine native/ Isabela |
| OM363354 | Ph_Native | Philippine native/ Isabela |
| OM363355 | Ph_Native | Philippine native/ Isabela |
| OM363356 | Ph_Native | Philippine native/ Isabela |
| OM363357 | Ph_Native | Philippine native/ Isabela |
| OM363358 | Ph_Native | Philippine native/ Isabela |
| OM363359 | Ph_Native | Philippine native/ Isabela |
| OM363360 | Ph_Native | Philippine native/ Isabela |
| OM363361 | Ph_Native | Philippine native/ Isabela |
| OM363362 | Ph_Native | Philippine native/ Isabela |
| OM363363 | Ph_Native | Philippine native/ Isabela |
| OM363364 | Ph_Native | Philippine native/ Isabela |
| OM363365 | Ph_Native | Philippine native/ Isabela |
| OM363366 | Ph_Native | Philippine native/ Isabela |
| OM363367 | Ph_Native | Philippine native/ Kalinga |
| OM363368 | Ph_Native | Philippine native/ Kalinga |
| OM363370 | Ph_Native | Philippine native/ Kalinga |
| OM363371 | Ph_Native | Philippine native/ Kalinga |
| OM363372 | Ph_Native | Philippine native/ Kalinga |
| OM363373 | Ph_Native | Philippine native/ Kalinga |
| OM363374 | Ph_Native | Philippine native/ Kalinga |
| OM363375 | Ph_Native | Philippine native/ Kalinga |
| OM363376 | Ph_Native | Philippine native/ Kalinga |
| OM363377 | Ph_Native | Philippine native/ Kalinga |
| OM363378 | Ph_Native | Philippine native/ Kalinga |
| OM363379 | Ph_Native | Philippine native/ Kalinga |
| OM363380 | Ph_Native | Philippine native/ Kalinga |
| OM363381 | Ph_Native | Philippine native/ Kalinga |
| OM363382 | Ph_Native | Philippine native/ Kalinga |
| OM363383 | Ph_Native | Philippine native/ Kalinga |
| OM363385 | Ph_Native | Philippine native/ Kalinga |
| OM363386 | Ph_Native | Philippine native/ Kalinga |
| OM363387 | Ph_Native | Philippine native/ Kalinga |
| OM363388 | Ph_Native | Philippine native/ Kalinga |
| OM363389 | Ph_Native | Philippine native/ Kalinga |
| OM363390 | Ph_Native | Philippine native/ Kalinga |
| OM363391 | Ph_Native | Philippine native/ Kalinga |
| OM363392 | Ph_Native | Philippine native/ Kalinga |
| OM363393 | Ph_Native | Philippine native/ Kalinga |
| OM363394 | Ph_Native | Philippine native/ Kalinga |
| OM363369 | Ph_Native | Philippine native/ Kalinga (interspecific hybrid of *S. philippensis x scrofa)* |
| OM363395 | Ph_Native | Philippine native/ Marinduque |
| OM363396 | Ph_Native | Philippine native/ Marinduque |
| OM363397 | Ph_Native | Philippine native/ Marinduque |
| OM363398 | Ph_Native | Philippine native/ Marinduque |
| OM363399 | Ph_Native | Philippine native/ Marinduque |
| OM363400 | Ph_Native | Philippine native/ Marinduque |
| OM363401 | Ph_Native | Philippine native/ Marinduque |
| OM363402 | Ph_Native | Philippine native/ Marinduque |
| OM363403 | Ph_Native | Philippine native/ Marinduque |
| OM363404 | Ph_Native | Philippine native/ Marinduque |
| OM363405 | Ph_Native | Philippine native/ Marinduque |
| OM363406 | Ph_Native | Philippine native/ Marinduque |
| OM363407 | Ph_Native | Philippine native/ Marinduque |
| OM363408 | Ph_Native | Philippine native/ Marinduque |
| OM363409 | Ph_Native | Philippine native/ Marinduque |
| OM363410 | Ph_Native | Philippine native/ Marinduque |
| OM363411 | Ph_Native | Philippine native/ Marinduque |
| OM363412 | Ph_Native | Philippine native/ Marinduque |
| OM363413 | Ph_Native | Philippine native/ Marinduque |
| OM363414 | Ph_Native | Philippine native/ Nueva Viscaya |
| OM363415 | Ph_Native | Philippine native/ Nueva Viscaya |
| OM363416 | Ph_Native | Philippine native/ Nueva Viscaya |
| OM363417 | Ph_Native | Philippine native/ Nueva Viscaya |
| OM363419 | Ph_Native | Philippine native/ Nueva Viscaya |
| OM363420 | Ph_Native | Philippine native/ Nueva Viscaya |
| OM363421 | Ph_Native | Philippine native/ Nueva Viscaya |
| OM363422 | Ph_Native | Philippine native/ Nueva Viscaya |
| OM363423 | Ph_Native | Philippine native/ Nueva Viscaya |
| OM363424 | Ph_Native | Philippine native/ Nueva Viscaya |
| OM363425 | Ph_Native | Philippine native/ Nueva Viscaya |
| OM363426 | Ph_Native | Philippine native/ Nueva Viscaya |
| OM363427 | Ph_Native | Philippine native/ Nueva Viscaya |
| OM363428 | Ph_Native | Philippine native/ Nueva Viscaya |
| OM363429 | Ph_Native | Philippine native/ Nueva Viscaya |
| OM363430 | Ph_Native | Philippine native/ Nueva Viscaya |
| OM363431 | Ph_Native | Philippine native/ Nueva Viscaya |
| OM363432 | Ph_Native | Philippine native/ Nueva Viscaya |
| OM363433 | Ph_Native | Philippine native/ Nueva Viscaya |
| MN625805.1 | Ph_Native | Philippine native/ Panay |
| MN625806.1 | Ph_Native | Philippine native/ Panay |
| MN625807.1 | Ph_Native | Philippine native/ Panay |
| MN625808.1 | Ph_Native | Philippine native/ Panay |
| MN625809.1 | Ph_Native | Philippine native/ Panay |
| MN625810.1 | Ph_Native | Philippine native/ Panay |
| MN625811.1 | Ph_Native | Philippine native/ Panay |
| MN625812.1 | Ph_Native | Philippine native/ Panay |
| MN625813.1 | Ph_Native | Philippine native/ Panay |
| MN625814.1 | Ph_Native | Philippine native/ Panay |
| MN625815.1 | Ph_Native | Philippine native/ Panay |
| MN625816.1 | Ph_Native | Philippine native/ Panay |
| MN625817.1 | Ph_Native | Philippine native/ Panay |
| MN625818.1 | Ph_Native | Philippine native/ Panay |
| MN625819.1 | Ph_Native | Philippine native/ Panay |
| MN625820.1 | Ph_Native | Philippine native/ Panay |
| MN625821.1 | Ph_Native | Philippine native/ Panay |
| MN625822.1 | Ph_Native | Philippine native/ Panay |
| MN625823.1 | Ph_Native | Philippine native/ Panay |
| MN625824.1 | Ph_Native | Philippine native/ Panay |
| MN625825.1 | Ph_Native | Philippine native/ Panay |
| MN625826.1 | Ph_Native | Philippine native/ Panay |
| MN625827.1 | Ph_Native | Philippine native/ Panay |
| MN625828.1 | Ph_Native | Philippine native/ Panay |
| OM363267 | Ph_Native | Philippine native/ Quezon |
| OM363434 | Ph_Native | Philippine native/ Quezon |
| OM363435 | Ph_Native | Philippine native/ Quezon |
| OM363436 | Ph_Native | Philippine native/ Quezon |
| OM363437 | Ph_Native | Philippine native/ Quezon |
| OM363438 | Ph_Native | Philippine native/ Quezon |
| OM363439 | Ph_Native | Philippine native/ Quezon |
| OM363440 | Ph_Native | Philippine native/ Quezon |
| OM363441 | Ph_Native | Philippine native/ Quezon |
| OM363442 | Ph_Native | Philippine native/ Quezon |
| OM363443 | Ph_Native | Philippine native/ Quezon |
| OM363444 | Ph_Native | Philippine native/ Quezon |
| OM363445 | Ph_Native | Philippine native/ Quezon |
| OM363446 | Ph_Native | Philippine native/ Quezon |
| OM363447 | Ph_Native | Philippine native/ Quezon |
| OM363448 | Ph_Native | Philippine native/ Quezon |
| OM363449 | Ph_Native | Philippine native/ Quezon |
| OM363450 | Ph_Native | Philippine native/ Quezon |
| OM363451 | Ph_Native | Philippine native/ Quezon |
| OM363452 | Ph_Native | Philippine native/ Quezon |
| OM363453 | Ph_Native | Philippine native/ Quezon |
| OM363454 | Ph_Native | Philippine native/ Quezon |
| OM363325 | Ph_Native | Philippine native/ Samar |
| OM363326 | Ph_Native | Philippine native/ Samar |
| OM363327 | Ph_Native | Philippine native/ Samar |
| OM363328 | Ph_Native | Philippine native/ Samar |
| OM363329 | Ph_Native | Philippine native/ Samar |
| OM363330 | Ph_Native | Philippine native/ Samar |
| OM363331 | Ph_Native | Philippine native/ Samar |
| OM363332 | Ph_Native | Philippine native/ Samar |
| OM363333 | Ph_Native | Philippine native/ Samar |
| OM363334 | Ph_Native | Philippine native/ Samar |
| OM363335 | Ph_Native | Philippine native/ Samar |
| OM363336 | Ph_Native | Philippine native/ Samar |
| OM363337 | Ph_Native | Philippine native/ Samar |
| OM363338 | Ph_Native | Philippine native/ Samar |
| OM363339 | Ph_Native | Philippine native/ Samar |
| OM363340 | Ph_Native | Philippine native/ Samar |
| OM363341 | Ph_Native | Philippine native/ Samar |
| OM363342 | Ph_Native | Philippine native/ Samar |
| OM363343 | Ph_Native | Philippine native/ Samar |
| OM363344 | Ph_Native | Philippine native/ Samar |
| AB041483.1 | Transboundary | Berkshire |
| AB041484.1 | Transboundary | Berkshire |
| AB041485.1 | Transboundary | Berkshire |
| AB059650.1 | Transboundary | Berkshire |
| AM040639.1 | Transboundary | Berkshire |
| AY429459.1 | Transboundary | Berkshire |
| AY574045.1 | Transboundary | Berkshire |
| AY884783.1 | Transboundary | Berkshire |
| DQ152897.2 | Transboundary | Berkshire |
| DQ152899.2 | Transboundary | Berkshire |
| DQ379191.2 | Transboundary | Berkshire |
| GQ169778.1 | Transboundary | Berkshire |
| KC505410.1 | Transboundary | Berkshire |
| KP765602.1 | Transboundary | Berkshire |
| OM363297 | Transboundary | Berkshire |
| OM363298 | Transboundary | Berkshire |
| OM363299 | Transboundary | Berkshire |
| AB041486.1 | Transboundary | Duroc |
| AB041487.2 | Transboundary | Duroc |
| AM040632.1 | Transboundary | Duroc |
| AY232880.1 | Transboundary | Duroc |
| AY232881.1 | Transboundary | Duroc |
| DQ379032.2 | Transboundary | Duroc |
| DQ379033.2 | Transboundary | Duroc |
| DQ379034.2 | Transboundary | Duroc |
| DQ379035.2 | Transboundary | Duroc |
| DQ379036.2 | Transboundary | Duroc |
| GQ141896.1 | Transboundary | Duroc |
| GQ141897.1 | Transboundary | Duroc |
| GQ169779.1 | Transboundary | Duroc |
| JX546493.1 | Transboundary | Duroc |
| JX546494.1 | Transboundary | Duroc |
| JX546495.1 | Transboundary | Duroc |
| JX546496.1 | Transboundary | Duroc |
| JX546497.1 | Transboundary | Duroc |
| KY964842.1 | Transboundary | Duroc |
| KY964843.1 | Transboundary | Duroc |
| KY964844.1 | Transboundary | Duroc |
| KY964845.1 | Transboundary | Duroc |
| KY964846.1 | Transboundary | Duroc |
| KY964847.1 | Transboundary | Duroc |
| KY964848.1 | Transboundary | Duroc |
| KY964849.1 | Transboundary | Duroc |
| KY964850.1 | Transboundary | Duroc |
| KY964851.1 | Transboundary | Duroc |
| OM363282 | Transboundary | Duroc |
| OM363283 | Transboundary | Duroc |
| OM363284 | Transboundary | Duroc |
| OM363285 | Transboundary | Duroc |
| OM363286 | Transboundary | Duroc |
| OM363287 | Transboundary | Duroc |
| OM363288 | Transboundary | Duroc |
| OM363289 | Transboundary | Duroc |
| AB041499.1 | Transboundary | Landrace |
| AY232884.1 | Transboundary | Landrace |
| AY232885.1 | Transboundary | Landrace |
| DQ379041.2 | Transboundary | Landrace |
| DQ379058.2 | Transboundary | Landrace |
| DQ379075.2 | Transboundary | Landrace |
| DQ379202.2 | Transboundary | Landrace |
| DQ379203.2 | Transboundary | Landrace |
| GQ141895.1 | Transboundary | Landrace |
| GQ169780.1 | Transboundary | Landrace |
| JX546512.1 | Transboundary | Landrace |
| JX546513.1 | Transboundary | Landrace |
| JX546514.1 | Transboundary | Landrace |
| JX546515.1 | Transboundary | Landrace |
| JX546516.1 | Transboundary | Landrace |
| KY964832.1 | Transboundary | Landrace |
| KY964833.1 | Transboundary | Landrace |
| KY964835.1 | Transboundary | Landrace |
| KY964836.1 | Transboundary | Landrace |
| KY964837.1 | Transboundary | Landrace |
| KY964838.1 | Transboundary | Landrace |
| KY964839.1 | Transboundary | Landrace |
| KY964840.1 | Transboundary | Landrace |
| KY964841.1 | Transboundary | Landrace |
| NC_000845.1 | Transboundary | Landrace |
| OM363275 | Transboundary | Landrace |
| OM363276 | Transboundary | Landrace |
| OM363277 | Transboundary | Landrace |
| OM363278 | Transboundary | Landrace |
| OM363279 | Transboundary | Landrace |
| OM363280 | Transboundary | Landrace |
| OM363281 | Transboundary | Landrace |
| AY230822.1 | Transboundary | Large White |
| AY232882.1 | Transboundary | Large White |
| AY232883.1 | Transboundary | Large White |
| DQ379125.2 | Transboundary | Large White |
| DQ379126.2 | Transboundary | Large White |
| DQ379127.2 | Transboundary | Large White |
| DQ379198.2 | Transboundary | Large White |
| DQ379214.2 | Transboundary | Large White |
| JX546562.1 | Transboundary | Large White |
| JX546563.1 | Transboundary | Large White |
| JX546564.1 | Transboundary | Large White |
| JX546565.1 | Transboundary | Large White |
| JX546566.1 | Transboundary | Large White |
| JX546567.1 | Transboundary | Large White |
| NC_012095 | Transboundary | Large White |
| OM363268 | Transboundary | Large White |
| OM363269 | Transboundary | Large White |
| OM363270 | Transboundary | Large White |
| OM363271 | Transboundary | Large White |
| OM363272 | Transboundary | Large White |
| OM363273 | Transboundary | Large White |
| OM363274 | Transboundary | Large White |
| GQ141893.1 | Transboundary | Large Yorkshire |
| GQ141894.1 | Transboundary | Large Yorkshire |
| AB041490.1 | Transboundary | Yorkshire |
| AB041491.1 | Transboundary | Yorkshire |
| AB041494.1 | Transboundary | Yorkshire |
| AM040633.1 | Transboundary | Yorkshire |
| AM040634.1 | Transboundary | Yorkshire |
| AM040635.1 | Transboundary | Yorkshire |
| AM040636.1 | Transboundary | Yorkshire |
| AM040638.1 | Transboundary | Yorkshire |
| AY574048.1 | Transboundary | Yorkshire |
| EF590197.1 | Transboundary | Yorkshire |
| GQ169777.1 | Transboundary | Yorkshire |
| JN601074.1 | Transboundary | Yorkshire |
| JN601075.1 | Transboundary | Yorkshire |
| KC250275.1 | Transboundary | Yorkshire |
| KY964834.1 | Transboundary | Yorkshire |
| KY964898.1 | Transboundary | Yorkshire |
| KY964899.1 | Transboundary | Yorkshire |
| KY964900.1 | Transboundary | Yorkshire |
| KY964901.1 | Transboundary | Yorkshire |
| KY964902.1 | Transboundary | Yorkshire |
| KY964903.1 | Transboundary | Yorkshire |
| KY964904.1 | Transboundary | Yorkshire |
| KY964905.1 | Transboundary | Yorkshire |
| KY964906.1 | Transboundary | Yorkshire |
| KY964907.1 | Transboundary | Yorkshire |
| NC_008830.1 | Wild | *Phacochoerus africanus* |
| NC_026992.1 | Wild | *Sus barbatus* |
| NC_023541.1 | Wild | *Sus cebifrons* |
| MN625829.1 | Wild | *Sus sp.* |
| MN625830.1 | Wild | *Sus sp.* |
| NC_023536.1 | Wild | *Sus verrucosus* |

**Online Resource 2** Sequence and position of the variable nucleotides in the 19 Philippine haplotypes

| **Populations/ Clade** | **Haplotype Number** | **Nucleotide Position and Sequence** | | | | | | | | | | | | | | | | | | | | | |
| --- | --- | --- | --- | --- | --- | --- | --- | --- | --- | --- | --- | --- | --- | --- | --- | --- | --- | --- | --- | --- | --- | --- | --- |
|  |  | 13 | 56 | 102 | 109 | 146 | 147 | 179 | 206 | 241 | 244 | 266 | 267 | 271 | 288 | 370 | 408 | 417 | 439 | 466 | 500 | 507 | 525 |
| **Cordillera clade** | Ph_1 | A | A | C | A | T | T | T | T | T | T | T | G | C | C | T | A | C | C | A | A | G | T |
|  | Ph_2 | A | A | A | A | T | T | T | T | T | T | T | G | C | C | T | A | C | T | A | A | G | T |
| **Asian clade (D2)** | Ph_3 | A | G | C | T | C | T | T | T | T | T | C | A | T | T | T | A | C | C | G | G | A | C |
|  | Ph_4 | A | G | C | T | C | T | T | T | T | T | C | A | T | T | T | A | C | C | G | G | A | T |
|  | Ph_5 | A | G | C | T | C | T | T | T | T | T | C | A | T | T | T | A | C | C | A | G | A | T |
|  | Ph_6 | A | G | C | T | C | T | T | C | T | T | C | A | T | T | T | A | C | C | A | G | A | T |
|  | Ph_7 | A | G | C | T | C | T | T | C | T | T | C | A | T | T | T | A | C | T | A | G | A | T |
|  | Ph_8 | A | G | C | T | C | T | T | T | T | C | C | A | T | T | C | A | T | C | A | G | A | T |
|  | Ph_9 | A | G | C | T | C | T | T | C | T | C | C | A | T | T | C | A | T | C | A | G | A | T |
|  | Ph_10 | A | G | C | T | C | T | T | T | T | C | T | A | T | T | T | A | T | C | A | G | A | C |
|  | Ph_11 | A | G | C | T | C | T | T | C | T | C | C | A | T | T | T | G | T | C | A | G | A | C |
| **Southeast Asian (D7/ MTSEA)** | Ph_12 | G | G | C | T | C | C | T | C | T | C | C | A | T | T | T | A | T | C | A | G | A | T |
|  | Ph_13 | G | G | C | T | C | T | C | C | T | C | C | A | T | T | T | A | T | C | A | G | A | T |
|  | Ph_14 | G | G | C | T | C | T | C | T | T | C | C | A | T | T | T | A | T | C | A | G | A | T |
|  | Ph_15 | A | G | C | T | C | C | C | C | T | C | C | A | T | T | T | A | T | C | G | G | A | T |
|  | Ph_16 | A | G | C | T | C | C | C | C | T | C | C | A | T | T | T | A | T | C | G | G | A | C |
|  | Ph_17 | A | G | C | T | C | C | C | C | T | C | C | A | C | T | T | A | T | C | G | G | A | C |
|  | Ph_18 | A | G | C | T | C | C | C | C | T | C | T | A | T | T | T | A | T | C | G | G | A | C |
|  | Ph_19 | A | G | C | T | T | C | C | C | C | C | C | A | T | T | T | A | C | C | G | G | A | C |

D2, D7, MTSEA are indicated based on pig domestication sites according to Larson et al. (2005), Tanaka et al. (2008), Layos et al. (2022a).

**Online Resource 3** Collection site and haplotype number of the Philippine native pig samples

| **Accession number** | **Haplotype Number** | **Haplogroup/ Cluster** | **Province** | **Region Number** | **Reference** |
| --- | --- | --- | --- | --- | --- |
| OM363316 | Ph_9 | AMC | Benguet | CAR | This study |
| OM363304 | Ph_9 | Asian Mix Cluster (AMC) | Benguet | CAR | This study |
| OM363306 | Ph_1 | CC | Benguet | CAR | This study |
| OM363308 | Ph_1 | CC | Benguet | CAR | This study |
| OM363310 | Ph_1 | CC | Benguet | CAR | This study |
| OM363311 | Ph_1 | CC | Benguet | CAR | This study |
| OM363312 | Ph_1 | CC | Benguet | CAR | This study |
| OM363313 | Ph_1 | CC | Benguet | CAR | This study |
| OM363315 | Ph_1 | CC | Benguet | CAR | This study |
| OM363321 | Ph_1 | CC | Benguet | CAR | This study |
| OM363323 | Ph_1 | CC | Benguet | CAR | This study |
| OM363303 | Ph_1 | Cordillera Cluster (CC) | Benguet | CAR | This study |
| OM363320 | Ph_10 | NLC | Benguet | CAR | This study |
| OM363322 | Ph_10 | NLC | Benguet | CAR | This study |
| OM363324 | Ph_10 | NLC | Benguet | CAR | This study |
| OM363369 | Ph_5 | AMC | Kalinga | CAR | This study |
| OM363383 | Ph_5 | AMC | Kalinga | CAR | This study |
| OM363371 | Ph_7 | AMC | Kalinga | CAR | This study |
| OM363380 | Ph_7 | AMC | Kalinga | CAR | This study |
| OM363367 | Ph_9 | AMC | Kalinga | CAR | This study |
| OM363382 | Ph_9 | AMC | Kalinga | CAR | This study |
| OM363388 | Ph_9 | AMC | Kalinga | CAR | This study |
| OM363389 | Ph_9 | AMC | Kalinga | CAR | This study |
| OM363390 | Ph_9 | AMC | Kalinga | CAR | This study |
| OM363391 | Ph_9 | AMC | Kalinga | CAR | This study |
| OM363445 | Ph_9 | AMC | Kalinga | CAR | This study |
| OM363373 | Ph_1 | CC | Kalinga | CAR | This study |
| OM363375 | Ph_1 | CC | Kalinga | CAR | This study |
| OM363376 | Ph_1 | CC | Kalinga | CAR | This study |
| OM363379 | Ph_1 | CC | Kalinga | CAR | This study |
| OM363384 | Ph_1 | CC | Kalinga | CAR | This study |
| OM363385 | Ph_1 | CC | Kalinga | CAR | This study |
| OM363392 | Ph_1 | CC | Kalinga | CAR | This study |
| OM363368 | Ph_10 | NLC | Kalinga | CAR | This study |
| OM363370 | Ph_10 | NLC | Kalinga | CAR | This study |
| OM363372 | Ph_10 | NLC | Kalinga | CAR | This study |
| OM363374 | Ph_10 | NLC | Kalinga | CAR | This study |
| OM363377 | Ph_10 | NLC | Kalinga | CAR | This study |
| OM363387 | Ph_10 | NLC | Kalinga | CAR | This study |
| OM363393 | Ph_10 | NLC | Kalinga | CAR | This study |
| OM363394 | Ph_10 | NLC | Kalinga | CAR | This study |
| OM363378 | Ph_16 | SLVC | Kalinga | CAR | This study |
| OM363386 | Ph_16 | SLVC | Kalinga | CAR | This study |
| OM363444 | Ph_16 | SLVC | Kalinga | CAR | This study |
| OM363381 | Ph_17 | SLVC | Kalinga | CAR | This study |
| OM363317 | Ph_1 | CC | Mt. Province | CAR | This study |
| OM363318 | Ph_1 | CC | Mt. Province | CAR | This study |
| OM363319 | Ph_1 | CC | Mt. Province | CAR | This study |
| OM363307 | Ph_10 | NLC | Mt. Province | CAR | This study |
| OM363309 | Ph_10 | NLC | Mt. Province | CAR | This study |
| OM363314 | Ph_10 | NLC | Mt. Province | CAR | This study |
| OM363305 | Ph_10 | North Luzon Cluster (NLC) | Mt. Province | CAR | This study |
| OM363332 | Ph_16 | SLVC | Leyte | 8 | This study |
| OM363333 | Ph_16 | SLVC | Leyte | 8 | This study |
| OM363337 | Ph_4 | AMC | Samar | 8 | This study |
| OM363339 | Ph_4 | AMC | Samar | 8 | This study |
| OM363326 | Ph_6 | AMC | Samar | 8 | This study |
| OM363344 | Ph_6 | AMC | Samar | 8 | This study |
| OM363327 | Ph_12 | SLVC | Samar | 8 | This study |
| OM363331 | Ph_12 | SLVC | Samar | 8 | This study |
| OM363335 | Ph_12 | SLVC | Samar | 8 | This study |
| OM363336 | Ph_12 | SLVC | Samar | 8 | This study |
| OM363340 | Ph_12 | SLVC | Samar | 8 | This study |
| OM363341 | Ph_12 | SLVC | Samar | 8 | This study |
| OM363329 | Ph_13 | SLVC | Samar | 8 | This study |
| OM363328 | Ph_16 | SLVC | Samar | 8 | This study |
| OM363330 | Ph_16 | SLVC | Samar | 8 | This study |
| OM363338 | Ph_16 | SLVC | Samar | 8 | This study |
| OM363342 | Ph_16 | SLVC | Samar | 8 | This study |
| OM363343 | Ph_17 | SLVC | Samar | 8 | This study |
| OM363334 | Ph_19 | SLVC | Samar | 8 | This study |
| OM363325 | Ph_14 | South Luzon Visayas Cluster (SLVC) | Samar | 8 | This study |
| MN625808.1 | Ph_4 | AMC | Panay | 6 | Layos, et al. 2025 |
| MN625813.1 | Ph_4 | AMC | Panay | 6 | Layos, et al. 2027 |
| MN625806.1 | Ph_7 | AMC | Panay | 6 | Layos, et al. 2023 |
| MN625807.1 | Ph_9 | AMC | Panay | 6 | Layos, et al. 2024 |
| MN625825.1 | Ph_11 | NLC | Panay | 6 | Layos, et al. 2028 |
| MN625810.1 | Ph_13 | SLVC | Panay | 6 | Layos, et al. 2026 |
| MN625805.1 | Ph_16 | SLVC | Panay | 6 | Layos, et al. 2022 |
| OM363413 | Ph_4 | AMC | Marinduque | 4B | This study |
| OM363395 | Ph_9 | AMC | Marinduque | 4B | This study |
| OM363397 | Ph_9 | AMC | Marinduque | 4B | This study |
| OM363398 | Ph_9 | AMC | Marinduque | 4B | This study |
| OM363400 | Ph_9 | AMC | Marinduque | 4B | This study |
| OM363402 | Ph_9 | AMC | Marinduque | 4B | This study |
| OM363407 | Ph_9 | AMC | Marinduque | 4B | This study |
| OM363409 | Ph_9 | AMC | Marinduque | 4B | This study |
| OM363410 | Ph_9 | AMC | Marinduque | 4B | This study |
| OM363406 | Ph_10 | NLC | Marinduque | 4B | This study |
| OM363401 | Ph_15 | SLVC | Marinduque | 4B | This study |
| OM363408 | Ph_15 | SLVC | Marinduque | 4B | This study |
| OM363411 | Ph_15 | SLVC | Marinduque | 4B | This study |
| OM363412 | Ph_15 | SLVC | Marinduque | 4B | This study |
| OM363399 | Ph_16 | SLVC | Marinduque | 4B | This study |
| OM363403 | Ph_16 | SLVC | Marinduque | 4B | This study |
| OM363404 | Ph_16 | SLVC | Marinduque | 4B | This study |
| OM363405 | Ph_17 | SLVC | Marinduque | 4B | This study |
| OM363396 | Ph_18 | SLVC | Marinduque | 4B | This study |
| OM363447 | Ph_1 | CC | Quezon | 4A | This study |
| OM363453 | Ph_1 | CC | Quezon | 4A | This study |
| OM363439 | Ph_13 | SLVC | Quezon | 4A | This study |
| OM363448 | Ph_15 | SLVC | Quezon | 4A | This study |
| OM363434 | Ph_16 | SLVC | Quezon | 4A | This study |
| OM363435 | Ph_16 | SLVC | Quezon | 4A | This study |
| OM363436 | Ph_16 | SLVC | Quezon | 4A | This study |
| OM363437 | Ph_16 | SLVC | Quezon | 4A | This study |
| OM363438 | Ph_16 | SLVC | Quezon | 4A | This study |
| OM363440 | Ph_16 | SLVC | Quezon | 4A | This study |
| OM363441 | Ph_16 | SLVC | Quezon | 4A | This study |
| OM363442 | Ph_16 | SLVC | Quezon | 4A | This study |
| OM363443 | Ph_16 | SLVC | Quezon | 4A | This study |
| OM363446 | Ph_16 | SLVC | Quezon | 4A | This study |
| OM363449 | Ph_16 | SLVC | Quezon | 4A | This study |
| OM363450 | Ph_16 | SLVC | Quezon | 4A | This study |
| OM363451 | Ph_16 | SLVC | Quezon | 4A | This study |
| OM363452 | Ph_16 | SLVC | Quezon | 4A | This study |
| OM363454 | Ph_16 | SLVC | Quezon | 4A | This study |
| KP987306.1 | Ph_1 | CC | Batanes | 2 | Li et al., 2017 |
| KP987307.1 | Ph_2 | CC | Batanes | 2 | Li et al., 2017 |
| OM363345 | Ph_4 | AMC | Isabela | 2 | This study |
| OM363346 | Ph_4 | AMC | Isabela | 2 | This study |
| OM363347 | Ph_4 | AMC | Isabela | 2 | This study |
| OM363354 | Ph_4 | AMC | Isabela | 2 | This study |
| OM363355 | Ph_5 | AMC | Isabela | 2 | This study |
| OM363356 | Ph_5 | AMC | Isabela | 2 | This study |
| OM363357 | Ph_5 | AMC | Isabela | 2 | This study |
| OM363359 | Ph_5 | AMC | Isabela | 2 | This study |
| OM363363 | Ph_5 | AMC | Isabela | 2 | This study |
| OM363364 | Ph_5 | AMC | Isabela | 2 | This study |
| OM363366 | Ph_5 | AMC | Isabela | 2 | This study |
| OM363351 | Ph_7 | AMC | Isabela | 2 | This study |
| OM363365 | Ph_7 | AMC | Isabela | 2 | This study |
| OM363348 | Ph_9 | AMC | Isabela | 2 | This study |
| OM363358 | Ph_9 | AMC | Isabela | 2 | This study |
| OM363360 | Ph_9 | AMC | Isabela | 2 | This study |
| OM363350 | Ph_1 | CC | Isabela | 2 | This study |
| OM363361 | Ph_1 | CC | Isabela | 2 | This study |
| OM363362 | Ph_1 | CC | Isabela | 2 | This study |
| OM363349 | Ph_10 | NLC | Isabela | 2 | This study |
| OM363353 | Ph_10 | NLC | Isabela | 2 | This study |
| OM363352 | Ph_16 | SLVC | Isabela | 2 | This study |
| OM363424 | Ph_5 | AMC | Nueva Viscaya | 2 | This study |
| OM363432 | Ph_7 | AMC | Nueva Viscaya | 2 | This study |
| OM363429 | Ph_8 | AMC | Nueva Viscaya | 2 | This study |
| OM363415 | Ph_1 | CC | Nueva Viscaya | 2 | This study |
| OM363416 | Ph_1 | CC | Nueva Viscaya | 2 | This study |
| OM363419 | Ph_1 | CC | Nueva Viscaya | 2 | This study |
| OM363420 | Ph_1 | CC | Nueva Viscaya | 2 | This study |
| OM363421 | Ph_1 | CC | Nueva Viscaya | 2 | This study |
| OM363422 | Ph_1 | CC | Nueva Viscaya | 2 | This study |
| OM363423 | Ph_1 | CC | Nueva Viscaya | 2 | This study |
| OM363425 | Ph_1 | CC | Nueva Viscaya | 2 | This study |
| OM363426 | Ph_1 | CC | Nueva Viscaya | 2 | This study |
| OM363430 | Ph_1 | CC | Nueva Viscaya | 2 | This study |
| OM363433 | Ph_1 | CC | Nueva Viscaya | 2 | This study |
| OM363427 | Ph_11 | NLC | Nueva Viscaya | 2 | This study |
| OM363414 | Ph_15 | SLVC | Nueva Viscaya | 2 | This study |
| OM363418 | Ph_15 | SLVC | Nueva Viscaya | 2 | This study |
| OM363428 | Ph_3 | AMC | Quirino | 2 | This study |
| OM363431 | Ph_3 | AMC | Quirino | 2 | This study |
| OM363417 | Ph_1 | CC | Quirino | 2 | This study |
| OM363267 | - | European mtDNA | Quezon | - | This study |

**Online Resource 4** Haplotype diversity and gene flow in the Philippine native pigs

|  | **Genetic diversity measures** | | | | | | | | | | | | | | |
| --- | --- | --- | --- | --- | --- | --- | --- | --- | --- | --- | --- | --- | --- | --- | --- |
| **Population** | **n** | **S** | **h** | **Hd** | **Var Hd** | **Pi** | **ThetaNuc** | **k** | **Theta G** | **Tajima D** | **FuLi D*** | **FuLi F*** | **Fu Fs** | **Gst** | **Nm** |
| ***Ph Native*** | ***175*** | ***22*** | ***19*** | ***0.894*** | ***0.000*** | ***0.0130*** | ***0.0071*** | ***6.981*** | ***3.833*** | ***2.261**** | ***-1.423*** | ***-0.442***** | ***2.044*** | ***0.078*** | ***2.96*** |
| Benguet | 22 | 14 | 3 | 0.567 | 0.006 | 0.0112 | 0.0071 | 6.039 | 3.841 | 2.047* | -2.355** | -2.203** | 9.518 |  |  |
| Kalinga | 30 | 19 | 8 | 0.851 | 0.001 | 0.0126 | 0.0089 | 6.786 | 4.796 | 1.438 | -2.471** | -2.239* | 3.459 |  |  |
| Isabela | 22 | 18 | 7 | 0.848 | 0.002 | 0.0089 | 0.0092 | 4.805 | 4.938 | -0.099 | -2.355 | -2.203 | 1.799 |  |  |
| Nueva Viscaya | 20 | 20 | 8 | 0.832 | 0.003 | 0.0135 | 0.0105 | 7.285 | 5.637 | 1.11 | -2.232 | -2.094 | 2.101 |  |  |
| Quezon | 19 | 18 | 5 | 0.405 | 0.020 | 0.0072 | 0.0097 | 3.895 | 5.233 | -0.987 | -2.383 | -2.305 | 2.587 |  |  |
| Marinduque | 19 | 10 | 7 | 0.784 | 0.006 | 0.0061 | 0.0053 | 3.263 | 2.861 | 0.498 | -2.457 | -2.371 | 0.141 |  |  |
| Samar | 20 | 11 | 8 | 0.832 | 0.003 | 0.0071 | 0.0058 | 3.805 | 3.101 | 0.806 | -2.232 | -2.094 | -0.109 |  |  |

*n-number of sequences, S-number of segregating sites, h-number of haplotypes, Hd-haplotype diversity, VarHd-variance of Hd, Pi-nucleotide diversity (Pi), K-average number of differences, G+Cn)-GC content non-coding, Gst-coefficient of gene differentiation, Nm-gene flow.

**Online Resource 5** Philippine collaborators that have provided assistance for the collection of pig samples

| **Name** | **Institution** |
| --- | --- |
| Rene C. Santiago, DVM, Ph.D. | ^1^NSPRDC, BAI |
| Vea Roven E. Arellano | NSPRDC, BAI |
| Rico M. Panaligan | NSPRDC, BAI |
| Marcelino G. Saliw-an, M.Sc. | ^2^KSU |
| Sharmaine D. Codiam | KSU |
| Sonwrigth B. Maddul, Ph.D. | ^3^BSU |
| Madeline S. Kingan, Ph.D. | BSU |
| Justine P. Ayomen | BSU |
| Diadem Freiah Lumerio, M.Sc. | BSU |
| Karina Marie G. Nicolas, DVM, Ph.D. | ^4^ISU |
| Dorothy P. Pagbilao, DVM, PhD | ^5^NVSU |
| Kayvin Petipet, DVM | NVSU |
| Arnolfo M. Monleon, Ph.D. | ^6^MSC |
| Adelina Idanan | MSC |
| Felix A. Afable, Ph.D. | ^7^ESSU |
| Johanna C. Casillano, | ESSU |
| Rhea Palma A. Ortego, M.Sc. | ESSU |
| Anseline Jane R. Mitra | ^8^UPLB |
| Mary Boneth T. Fallena | UPLB |
| Elsie Erica C. Abes | UPLB |
| Katrina U. Aquino, M.Sc. | UPLB |
| Carla Alilie L. Junsay, M.Sc. | UPLB |
| Ana Clarissa M. Ambagan | UPLB |
| Medino Gedeun N. Yebron, Jr., M.Sc. | UPLB |
| Elpidio M. Agbisit, Jr., Ph.D. | UPLB |
| ^1^National Swine and Poultry Research and Development Center (NSPRDC), Bureau of Animal Industry, Tiaong Stock Farm, Tiaong, Quezon,  ^2^Kalinga State University (KSU), Tabuk City, Kalinga  ^3^Benguet State University (BSU), La Trinidad, Benguet  ^4^Isabela State University (ISU), Echague, Isabela  ^5^Nueva Viscaya State University (NVSU), Bagabag, Nueva Viscaya  ^6^Marinduque State College (MSC), Torrijos, Marinduque  ^7^Eastern Samar State University (ESSU), Borongan City, Eastern Samar  ^8^University of the Philippines Los Baños (UPLB), College, Laguna | |
